# Global biogeographical regions of freshwater fish species

**DOI:** 10.1101/319566

**Authors:** Boris Leroy, Murilo S. Dias, Emilien Giraud, Bernard Hugueny, Céline Jézéquel, Fabien Leprieur, Thierry Oberdorff, Pablo A. Tedesco

## Abstract

**Aim:** To define the major biogeographical regions and transition zones for freshwater fish species.

**Taxon:** Strictly freshwater species of actinopterygian fish (i.e., excluding marine and amphidromous fish families).

**Methods:** We based our bioregionalisation on a global database of freshwater fish species occurrences in drainage basins, which, after filtering, includes 11 295 species in 2 581 basins. On the basis of this dataset, we generated a bipartite (basin-species) network upon which we applied a hierarchical clustering algorithm (the Map Equation) to detect regions. We tested the robustness of regions with a sensitivity analysis. We identified transition zones between major regions with the participation coefficient, indicating the degree to which a basin has species from multiple regions.

**Results:** Our bioregionalisation scheme showed two major supercontinental regions (Old World and New World, 50% species of the world and 99.96% endemics each). Nested within these two supercontinental regions lie six major regions (Nearctic, Neotropical, Palearctic, Ethiopian, Sino-Oriental and Australian) with extremely high degrees of endemism (above 96% except for the Palearctic). Transition zones between regions were of limited extent compared to other groups of organisms. We identified numerous subregions with high diversity and endemism in tropical areas (e.g. Neotropical), and a few large subregions with low diversity and endemism at high latitudes (e.g. Palearctic).

**Main conclusions:** Our results suggest that regions of freshwater fish species were shaped by events of vicariance and geodispersal which were similar to other groups, but with freshwater-specific processes of isolation that led to extremely high degrees of endemism (far exceeding endemism rates of other continental vertebrates), specific boundary locations, and limited extents of transition zones. The identified bioregions and transition zones of freshwater fish species reflect the strong isolation of freshwater fish faunas for the past 10 to 20 million years. The extremely high endemism and diversity of freshwater fish fauna raises many questions about the biogeographical consequences of current introductions and extinctions.

## Introduction

For almost two centuries, biogeographers have classified continental areas of the world into distinct biogeographical regions on the basis of organism distributions across the Earth (Wallace, 1876; Cox, 2001; Holt et al., 2013). Indeed, early biogeographers observed that many organisms share constellated distributions of endemics in particular regions. Furthermore, they observed that these patterns of endemism are often similar for distinct groups of organisms, resulting in very similar biogeographical regions. This marked similarity has led to the hypothesis that these regions reflect a shared history of diversification among taxa and have been conditioned by geography, geology and climate (Morrone, 2015; Lomolino, Riddle, & Whittaker, 2016).

The earliest classifications outlined six major biogeographic regions for birds (Sclater, 1858) and non-flying mammals (Wallace, 1876) (Nearctic, Neotropical, Palearctic, Ethiopian, Oriental and Australian). During recent years, these major regions have been confirmed by an upsurge in bioregionalisation studies. This upsurge was facilitated by the increase in quality and quantity of large-scale datasets, as well as the development of new analytical tools (Kreft & Jetz, 2010; Vilhena & Antonelli, 2015; Edler, Guedes, Zizka, Rosvall, & Antonelli, 2016). Consequently, multiple studies have tried to identify the major biogeographical regions for birds (Procheş & Ramdhani, 2012; Rueda, Rodríguez, & Hawkins, 2013; Holt et al., 2013), mammals (Kreft & Jetz, 2010; Procheş & Ramdhani, 2012; Rueda et al., 2013; Holt et al., 2013), amphibians (Procheş & Ramdhani, 2012; Rueda et al., 2013; Holt et al., 2013; Vilhena & Antonelli, 2015; Edler et al., 2016) and reptiles (Procheş & Ramdhani, 2012). The result of this upsurge was a debate on the precise limits of biogeographical regions. Indeed, some studies explicitly defined transition zones as distinct regions (e.g., Holt et al., 2013), whereas others included transition zones in major regions (Kreft & Jetz, 2010). This question of transition zones was settled to some extent in the major synthesis of Morrone (2015), proposing that transition zones should not be considered as distinct regions, but rather as transitional boundaries between major regions. Indeed, some regions share sharp boundaries, reflecting a long history of isolation by tectonics (Ficetola, Mazel, & Thuiller, 2017), whereas others share diffuse boundaries, reflecting recent interchanges, generally limited by mountain or climatic barriers (Morrone, 2015; Ficetola et al., 2017). Morrone (2015) proposed that five major transition zones emerged from anterior works, which could be explained by a vicariance-dispersal model based on tectonic history. This synthetic model can be considered as a general framework to test for biogeographical regions.

However, the recent upsurge in continental bioregionalisation studies has concentrated exclusively on terrestrial vertebrates, which represent but a fraction of the continental organisms. There are other continental organisms with constraints to their dispersal and ecology that are fundamentally distinct from terrestrial vertebrates and for which one might expect distinct biogeographical regions. For example, terrestrial plants are generally characterised by higher degrees of endemism than animals, because they are more constrained than animals in terms of dispersal and tolerance to surmount climatic and other physical barriers (Lomolino et al., 2016). Hence, major phytogeographical regions were described as manifold small regions (De Candolle, 1820, 1855; Takhtajan, 1986). However, Cox (2001) later proposed a handful of large floral regions comparable to biogeographical regions, thus suggesting that the major biogeographical regions are universal across the tree of life. A second example concerns human microbial diseases whose biogeography has also been shown recently to match terrestrial vertebrate biogeography (Murray et al., 2015). Another possibility concerns strictly freshwater organisms (i.e., organisms that live and disperse exclusively in freshwaters) as they have lower dispersal abilities than terrestrial vertebrates, and are geographically isolated in drainage basins usually flowing to the oceans. Terrestrial boundaries and salt waters represent strong barriers to dispersal, hence drainage basins have been considered as ‘island-like’ systems for strictly freshwater organisms (Rahel, 2007; Hugueny, Oberdorff, & Tedesco, 2010; Tedesco et al., 2012; Dias et al., 2014). Dispersal can occur actively or passively via underground waters, stream captures, exceptional floods, glacier melting causing stream overflow, confluence during sea-level lowering, and displacement by other organisms or typhoons (see also discussion in Capobianco & Friedman, 2018). However, such dispersal events are rare, therefore immigration and speciation presumably occur on similar time-scales (Oberdorff et al., 2011). Consequently, one might expect that, because of peculiarities of riverscape changes through geological times, strictly freshwater organisms have been subject to different histories of diversification from those of terrestrial vertebrates (Rahel, 2007) and thus have original biogeographical boundaries. Because dispersal is physically constrained, a higher degree of provincialism and endemism could be anticipated for such organisms, resulting potentially in smaller and more numerous biogeographic regions.

In this paper, we focussed on the global biogeography of strictly freshwater actinopterygian fishes (i.e., excluding marine and amphidromous families of fish), hereafter called freshwater fishes. Several studies delineated biogeographical regions of freshwater fishes at regional to continental scales (e.g., Unmack 2001; Oikonomou *et al*. 2014), and studies conducted at the global scale also focussed on subregional provinces (ecoregions) based on a combination of data and expert decisions (Abell et al., 2008; Lévêque, Oberdorff, Paugy, Stiassny, & Tedesco, 2008). Only one work hinted at nine potential freshwater fish biogeographic regions that covered the same biogeographical regions as terrestrial vertebrates (Matthews, 1998), but this work was based on a coarse geographic scale (52 approximate drainage basins for the whole world) and a low taxonomic resolution (family level). In addition, Matthews, (1998) included marine and diadromous fish families, which could conceal the effect of long-term isolation on freshwater fish endemicity patterns. Consequently, whether Sclater-Wallace’s biogeographical regions are also applicable to freshwater fishes and to other freshwater organisms remains unresolved, and a global-scale quantitative bioregionalisation would represent an important step forward.

In this study, we aimed to define the major biogeographical regions for strictly freshwater fish species at the global scale. To delineate biogeographical regions, we capitalised on the recent development of a comprehensive dataset on freshwater fish distributions in drainage basins covering more than 80% of the Earth surface (Tedesco et al., 2017). First, we identified the large biogeographical regions of freshwater fishes using a recently developed hierarchical approach based on networks (Vilhena & Antonelli, 2015), recommended for bioregionalisation studies (Edler et al., 2016; Bloomfield, Knerr, & Encinas-Viso, 2017; Rojas, Patarroyo, Mao, Bengtson, & Kowalewski, 2017). Then, we mapped the transition zones between regions and investigated species distributed across region boundaries. Finally, we compared biogeographical regions with terrestrial vertebrate biogeographical regions and discussed our findings in light of the synthetic biogeographical model proposed by Morrone (2015).

## Methods

### Distribution data

We based our bioregionalisation on the most comprehensive global database on freshwater fish species occurrence in drainage basins (Tedesco et al., 2017). This database comprises 110 331 occurrence records for 14 953 species in 3 119 drainage basins of the world. Species names in the database were validated according to FishBase (Froese & Pauly, 2017) and the Catalogue of Fishes (Fricke, Eschmeyer, & van der Laan, 2017), and occurrence records were screened by the team developing the database (see details in Tedesco *et al*. 2017). We applied additional filters and corrections to the database. Since our aim was to describe the natural biogeographical regions resulting from long-term isolation of freshwater ichthyofaunas, we excluded documented records of introduced species, but included species considered to be recently extinct in their historical river basins. Additionally, to exclude most species that could disperse through marine waters, we retained only families having less than 10% of their species occurring in marine waters. This filter retained all “primary” and almost all “secondary” families of fishes (only Pseudomugilidae and Fundulidae were excluded), i.e. families with respectively no or limited salt tolerant species, as well as 22 families that had never been classified (based on Table 2 of Berra, 2007). It also included eight families with marine ancestors, seven of which had no species classified as tolerating salt water. Finally, we removed all diadromous species, according to FishBase. Additionally, we detected a few errors that were corrected in the database, mostly related to the native/introduced status for some species. The database used in this publication is available in Appendix S1 in Supporting Information. The resulting dataset included 59 373 records of 11 295 species in 2 581 basins (Figure 1).

**Figure 1.**
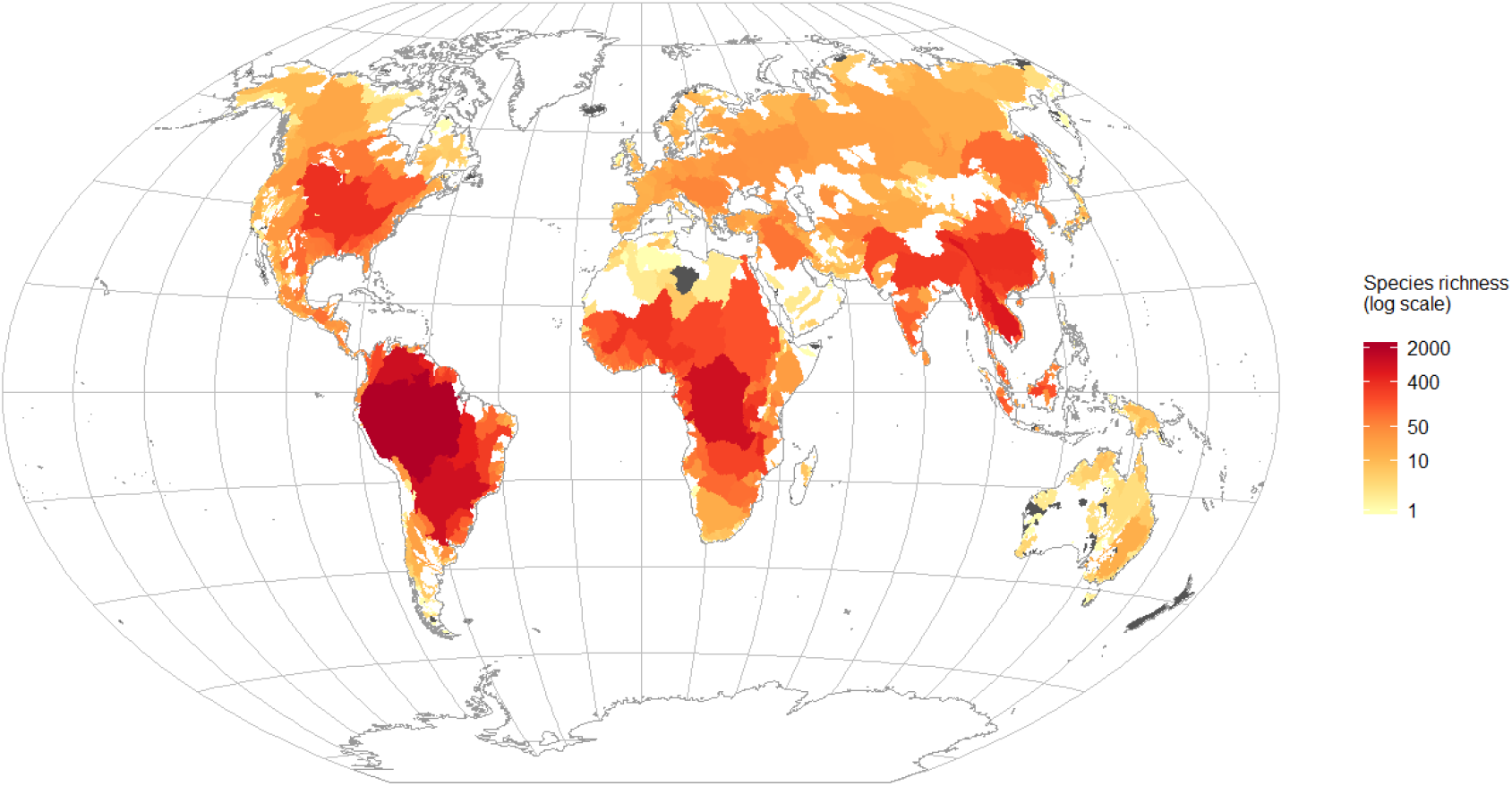
Global distribution of freshwater fish species richness per drainage basin based on the global database on freshwater fish species occurrence in drainage basins (Tedesco et al., 2017). Grey-shaded areas correspond to basins without records of native strictly freshwater species.

To define our bioregions, we worked at the species level and used drainage basins as geographical units. Indeed, (1) in the absence of a unified phylogeny for actinopterygian fishes, species is the most standard unit available and (2) contrary to terrestrial vertebrates, (for which gridded distribution data of reliable quality are is available), the most precise distribution data available for actinopterygian fishes is at the drainage basin unit. However, it is important to note that even if drainage basins are uneven in size, they are biogeographically meaningful for freshwater organisms because water bodies are generally connected within basins but not between basins (Hugueny et al., 2010).

### Delineation of biogeographical regions

Until recently, the prevailing procedure for bioregionalisation has been based on hierarchical clustering methods applied to compositional dissimilarity (Kreft & Jetz, 2010; Procheş & Ramdhani, 2012; Holt et al., 2013). Since then, an approach based on biogeographical networks was introduced by Vilhena & Antonelli (2015), and has been recommended for delineating biogeographic regions (Edler et al., 2016; Bloomfield et al., 2017; Rojas et al., 2017). A network is composed of a series of *nodes* which can be connected to each other by *links* (or edges). In bioregionalisation, the network is composed of both sites (i.e., drainage basins here) and species, which constitutes a *bipartite* network. When a taxon is known to occur at a particular site, a link is drawn between the taxon and the site. A site cannot be connected to another site, and a taxon cannot be connected to another taxon. By definition, site-site and species-species links are not allowed in this type of analysis. Our final network had 13 876 nodes (11 295 species and 2 581 basins) and 59 373 links. We handled the network under Gephi 0.9.2, with the ForceAtlas2 algorithm. This software groups nodes that are tightly interconnected (such as groups of sites and species from the same biogeographical region) and separates groups of nodes that are not interconnected (distinct biogeographical regions). Such a graphical representation is useful for analysing and exploring the network.

We applied a community-detection algorithm to the entire network in order to group nodes into clusters (i.e. biogeographical regions). We applied the Map Equation algorithm (www.mapequation.org, Rosvall & Bergstrom, 2008) because it has been tested and recommended to identify biogeographical regions (Vilhena & Antonelli, 2015; Edler et al., 2016; Rojas et al., 2017) and it features hierarchical clustering. Clusters are identified by the algorithm as having high intra-group but low inter-group connectivity, which corresponds well to the definition of biogeographical regions, i.e. regions of distinct assemblages of endemic taxa. We ran Map Equation (version Sat Oct 28 2017) with 100 trials to find the optimal clustering. We ran the hierarchical clustering (i.e., multi-level) in order to test whether larger regions have a nested hierarchy of subregions. It is important to note that a hierarchy of regions identified at the species level illustrates how biogeographical regions (i.e., distinct assemblages of endemic taxa) are currently spatially nested, but does not represent a historical (i.e., evolutionary) hierarchy of how these regions emerged.

The biogeographical network approach presents several advantages over distance-based approaches that were instrumental in our choice. Foremost, species identities are not lost, i.e., they are not abstracted into dissimilarity matrices between sites. Consequently, the network approach allows one to map how sites are connected by individual species, which presents an unquestionable asset to investigate between- and within-regions structures, such as potential dispersal pathways or barriers. A second practical novelty is that the algorithm assigns each species to a specific bioregion, which enables species-level descriptions (e.g. for online databases such as FishBase) and analyses. Lastly, the Map Equation algorithm is robust to differences in sampling intensities, making the removal of basins with low species richness unnecessary. On the other hand, distance-based approaches have limitations (see e.g., Leprieur & Oikonomou, 2014) and can produce inconsistent results when transforming such large occurrence datasets into a single dimension during the clustering procedure (see Appendix S2).

However, we provide clustering results using two additional methods in Appendix S2 for comparison: another network-based algorithm (Simulated Annealing, Bloomfield et al., 2017) and a distance-based method (following the framework of Kreft & Jetz, 2010).

### Sensitivity analysis

We analysed the robustness of the identified regions by randomly extirpating a percentage of species (random value between 0.01 and 10.00% of the total number of species in the database) and re-running the whole bioregionalisation process. This process was repeated 200 times. Then, for each region, we quantified the percentage of each region initial area that was retrieved in each simulation (Appendix S3).

### Transition zones and species shared between regions

We calculated the participation coefficient (Guimerà & Amaral, 2005; Bloomfield et al., 2017) for each node of the biogeographical network. The participation coefficient indicates the degree to which a node is connected to different regions. A high participation coefficient for a given basin indicates that it contains species from different regions and can be assimilated to a transition zone between regions. A low participation coefficient indicates that all species in the basin belong to the same region. The participation coefficient of a node is calculated as follows:

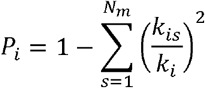

where *P_i_* is the participation coefficient of node *i, k_is_* is the number of links of node *i* to region *s, k_i_* is the total number of links of node *i*, and *N*_m_ is the total number of regions. We calculated the participation coefficient at each level of the biogeographical structure identified by Map Equation.

We also summarised the list of species that were shared between the major regions (i.e., excluding tiny clusters) and their distribution characteristics.

## Results

The Map Equation algorithm identified a hierarchy of biogeographical regions with up to six nested levels. For this global-scale study, we investigated the first three levels, termed as supercontinental regions, Regions and Subregions.

### Supercontinental regions

At the first level, we found that the world of freshwater fishes was divided into two supercontinental regions that we named New World (Americas) and Old World (Eurasia, Africa and Australian) (Figure 2a). Each supercontinental region contained nearly half the world’s 11 295 species with virtually 100% of endemic species (Table 1). Only two species occurred in both supercontinental regions (Figure 3 and Appendix S4): (1) *Esox lucius*, an Old World species that occurred in 32 basins of northern New World, and (2) *Catostomus catostomus*, a New World species that occurred in nine basins of northern Old World. These two supercontinental regions hosted 99% of endemic genera and around 80% of endemic families. At this first level, we also found 14 tiny clusters of 49 basins (exclusively located in the Old World) without endemic families and genera but with endemic species (40 species in total). These tiny clusters were most often composed of species-poor basins located in remote islands (e.g., Madagascan) or isolated arid areas (e.g., Arabian Peninsula). Therefore, we *post-hoc* assigned these clusters to the Old World supercontinental region (see details in Appendix S5).

**Figure 2.**
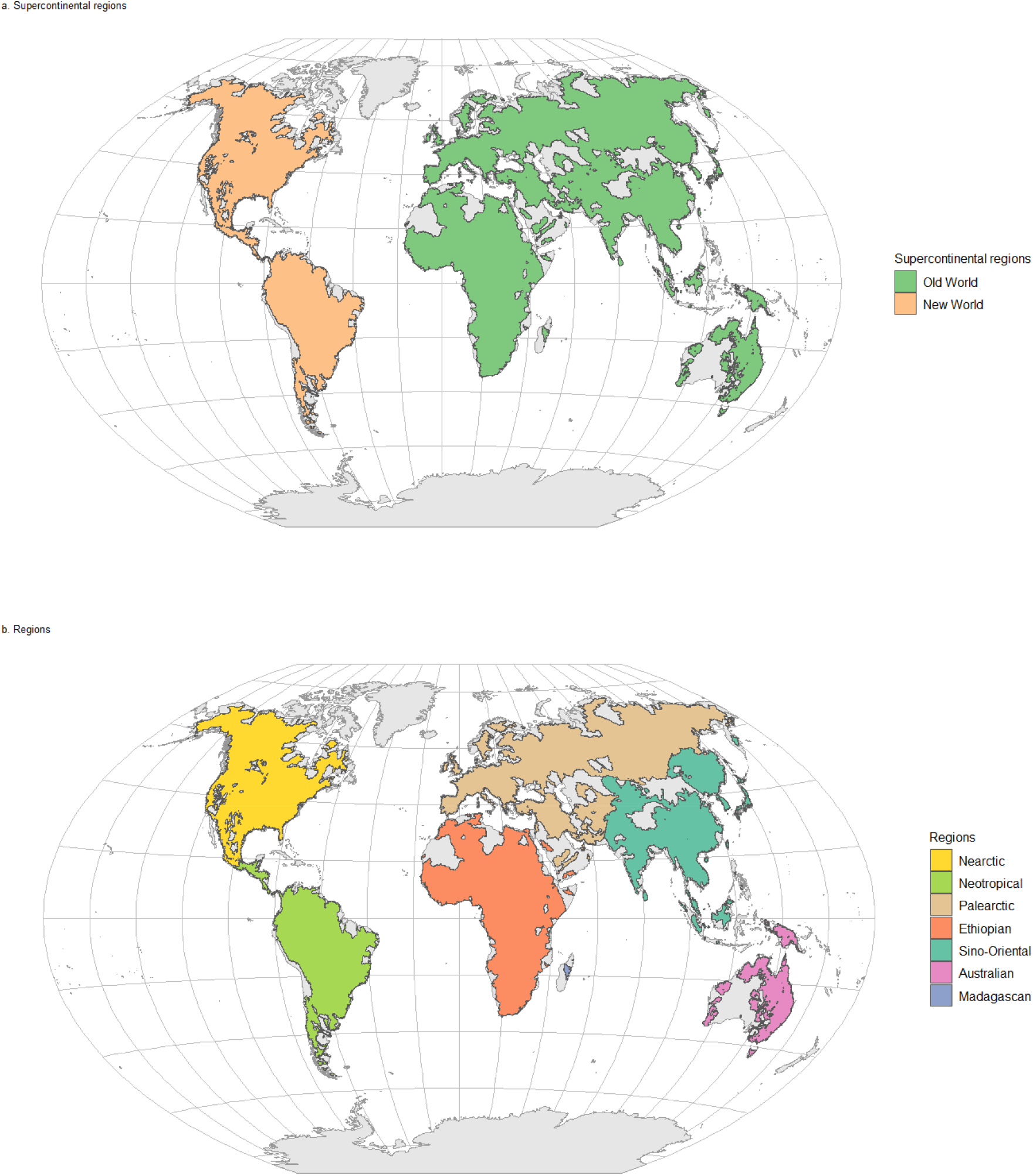
Biogeographical regions of freshwater fishes defined at the species level with the Map Equation clustering algorithm. We identified (a) two major supercontinental regions (Old World and New World) and (b) six major regions (Nearctic, Palearctic, Neotropical, Ethiopian, Sino-Oriental, Australian) and a minor cluster (Madagascan). Some very small clusters of a few drainage basins that do not share species with any other basin were corrected based on expert interpretation and literature (see details in Appendix S5).

**Figure 3.**
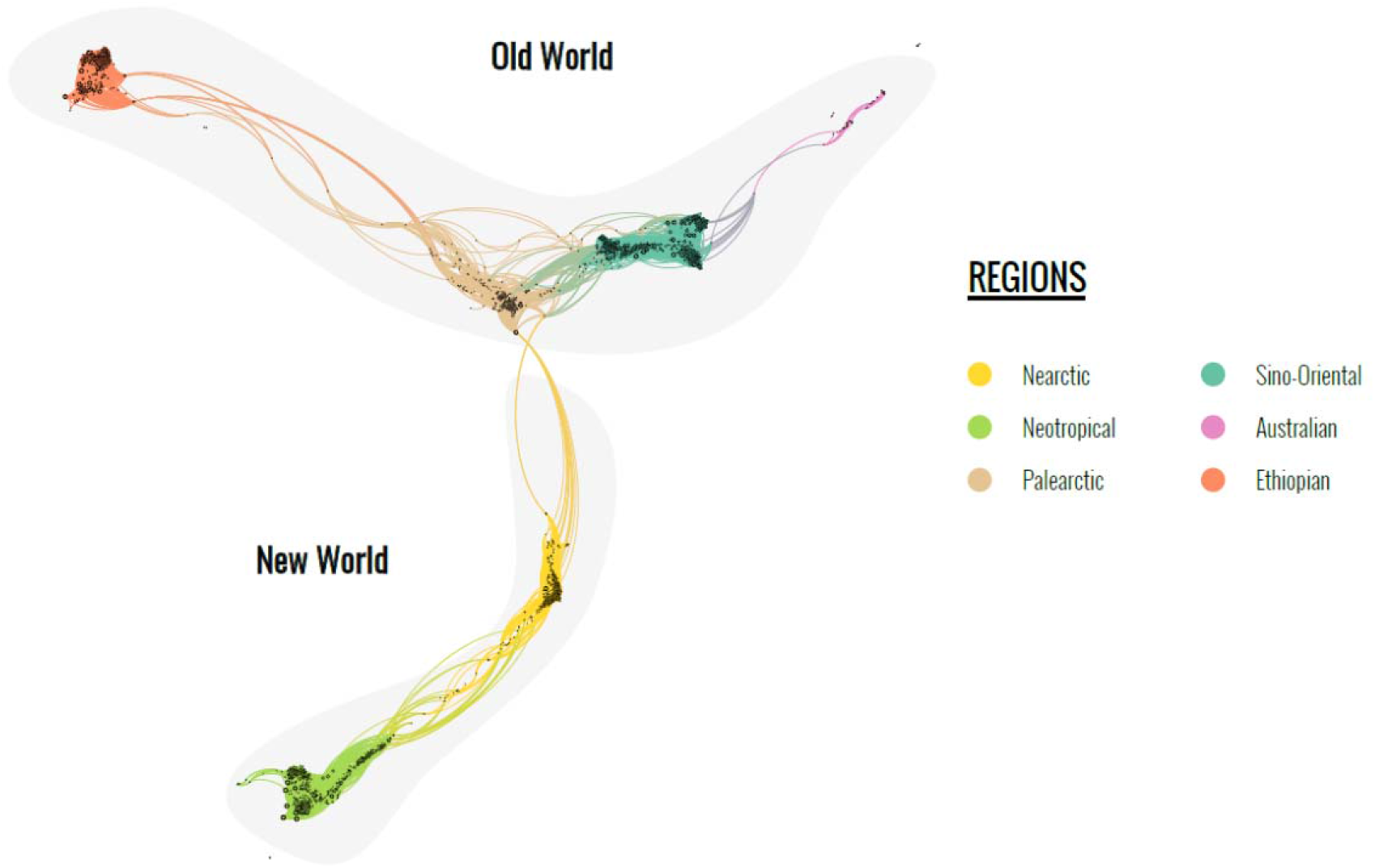
Global biogeographical network of freshwater fishes. In this network, both species and drainage basins are represented as nodes. When a species is known to occur in a drainage basin, a link between the species and the basin is drawn. The network is very complex because of the high number of nodes (13 876 nodes corresponding to 11 295 species and 2 581 basins) and links (59 373 occurrences). We spatialised the network in Gephi with the ForceAtlas 2 algorithm in order to group nodes that are strongly interconnected (i.e., basins that share species in common) and spread away from all other nodes that are not interconnected (i.e., basins that have few or no species in common). We coloured species and basin nodes according to the regions identified with the Map Equation algorithm and highlighted in grey the two supercontinental region (New World and Old World). To simplify the network, we masked here all nodes with less than ten links. A zoomable version of the full network with species and basin names is available in Appendix S6.

**Table 1.**
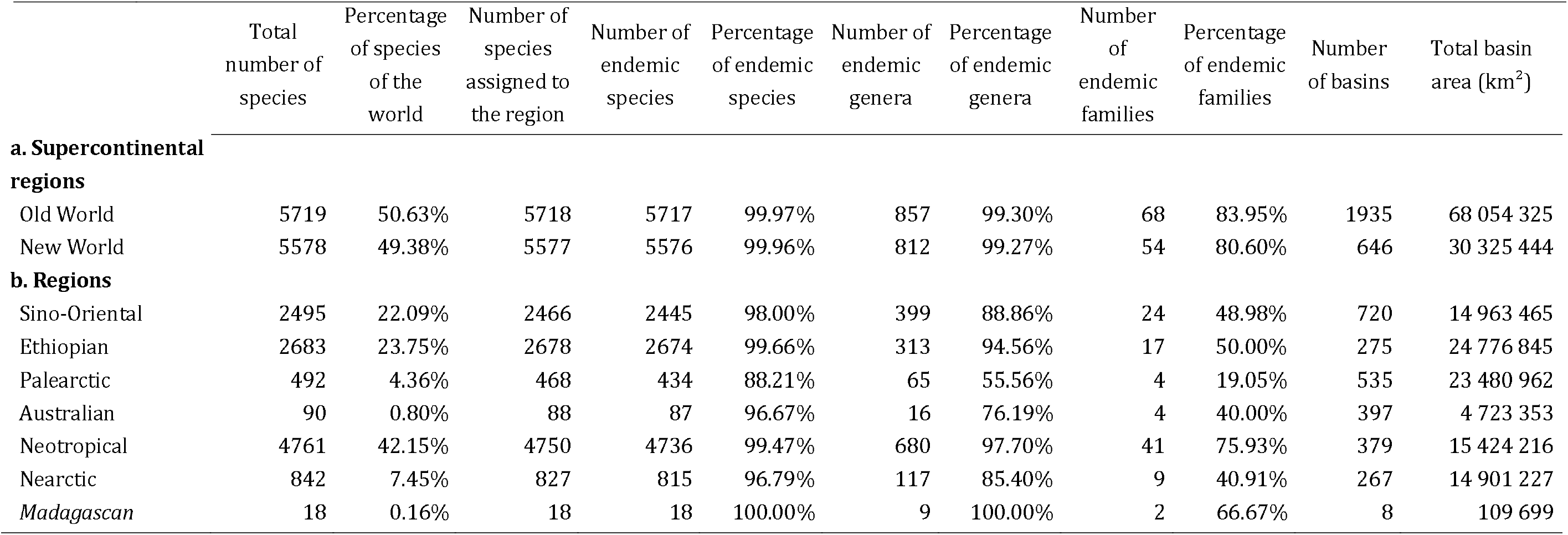
Characteristics of the first two levels of biogeographical regions of freshwater fishes identified with the Map Equation algorithm.

### Regions

At the second level, we found six major regions spatially nested within the two supercontinental regions (Figure 2b), that we named following Morrone (2015). In the Old World supercontinental region, we found four regions and a minor cluster (Figure 2b). The richest one (Table 1) was the Ethiopian region with nearly 50% of Old World species, covering the entire African continent and including areas north from the Sahara and a few basins in the Arabic peninsula. The second richest one was the Sino-Oriental region which included south-eastern Asia from India to Borneo, most of China and Mongolia, Korea and Japan. The third one was the Palearctic region with less than 10% of Old World species, covering Europe, Central Asia (up to Pakistan and Kazakhstan) and Siberia. The fourth one, the poorest in species, was the Australian region, with only 80 species in total, covering Australia, Tasmania and Papouasia-New Guinea. Last, we identified Madagascan as a distinct minor cluster of the Old World, with 100% of endemic species and genera. Within the New World supercontinental region, we found two major regions. The first one was the Neotropical region, containing 85% of New World species and 42% of the world’s known freshwater fish species (Table 1). The Neotropical region covered the whole of South America and Mesoamerica up to Southern Mexico. The second one was the Nearctic region, covering North America and northern Mexico. Finally, a tiny cluster composed of four basins of Central America was identified at this level, which we *post-hoc* assigned to the Nearctic region (see details in Appendix S5).

Most regions had very high degrees of endemism (Table 1b), above 96% for all regions except the Palearctic (88%). These high degrees of endemism are apparent on the biogeographical network through a low number of links between regions (Figure 3, Appendix S6). In other words, all regions shared only a very limited number of species (see Appendix S4). The degree of endemism was lower for genera, ranging from 56% for the Palearctic to 98% for the Neotropical, and was much lower for families, with values below or equal to 50% for all regions except the Neotropical one.

Interestingly, our clustering results are concordant with both Simulated Annealing (Figure S2.1 in Appendix S2) and beta-diversity methods (Figure S2.2). The only notable discrepancies concerned the Nearctic region which was split in four regions with the beta-diversity approach, and a portion of the Central Asian part of the Sino-Oriental which was attributed to the Palearctic by the Simulated Annealing method (see Appendix S2). Likewise, our sensitivity analysis on the Map Equation algorithm confirmed that regions were stable to random extirpations of species (Appendix S3), except for the Sino-Oriental one, which split in two regions in half of the simulations. This split produced the Oriental and Sinean regions with respectively ∼85% and ∼72% of endemics. These two subregions are visible on the network with two apparent clusters of nodes in the Sino-Oriental cluster (Figure 3). We also observed other minor changes, such as some clusters of basins appearing as small distinct regions, e.g. in Central Asia.

### Subregions

At the third level, we observed different patterns among regions. Three regions (Sino-Oriental, Nearctic and Australian) had only two to three main subregions (Appendix S7) that were spatially coherent and had high degrees of endemism (68.7-91.5% of endemic species, Appendix S7). The other three regions (Ethiopian, Palearctic and Neotropical) were characterised by a high number of subregions, which were also generally spatially coherent (Appendix S7). The Ethiopian and Neotropical subregions were generally characterised by a high number of species and endemics (Figures S7.9 and S7.14). The Palearctic subregions were characterised by a low number of species and generally low endemicity (Figure S7.11).

### Transition zones and species shared between regions

At the supercontinental region level, we obtained participation coefficients of basins between 0.0 and 0.5, with transition zones (i.e., basins with high participation coefficients) located in north-eastern Siberia as well as in northern North America. At the regional level, we observed participation coefficients also ranging from 0.0 to 0.5 (Figure 4). Unsurprisingly, we found the same transition zones between the Nearctic and the Palearctic as for supercontinental regions. However, the major transition zones were located at the boundaries between the Palearctic and Sino-Oriental regions: the high participation coefficients of basins at their boundaries indicated that these boundaries were diffuse (Figure 4). These diffuse boundaries are reflected on the network by the high number of links between multiple species and multiple basins from both regions (Figure 3, Appendix S5). We identified another transition zone between the Nearctic and the Neotropical regions, with high participation coefficients of basins at their boundary (Figure 4). A dozen species of each region also incurred in the other region with similar distribution patterns (Appendix S4).

**Figure 4.**
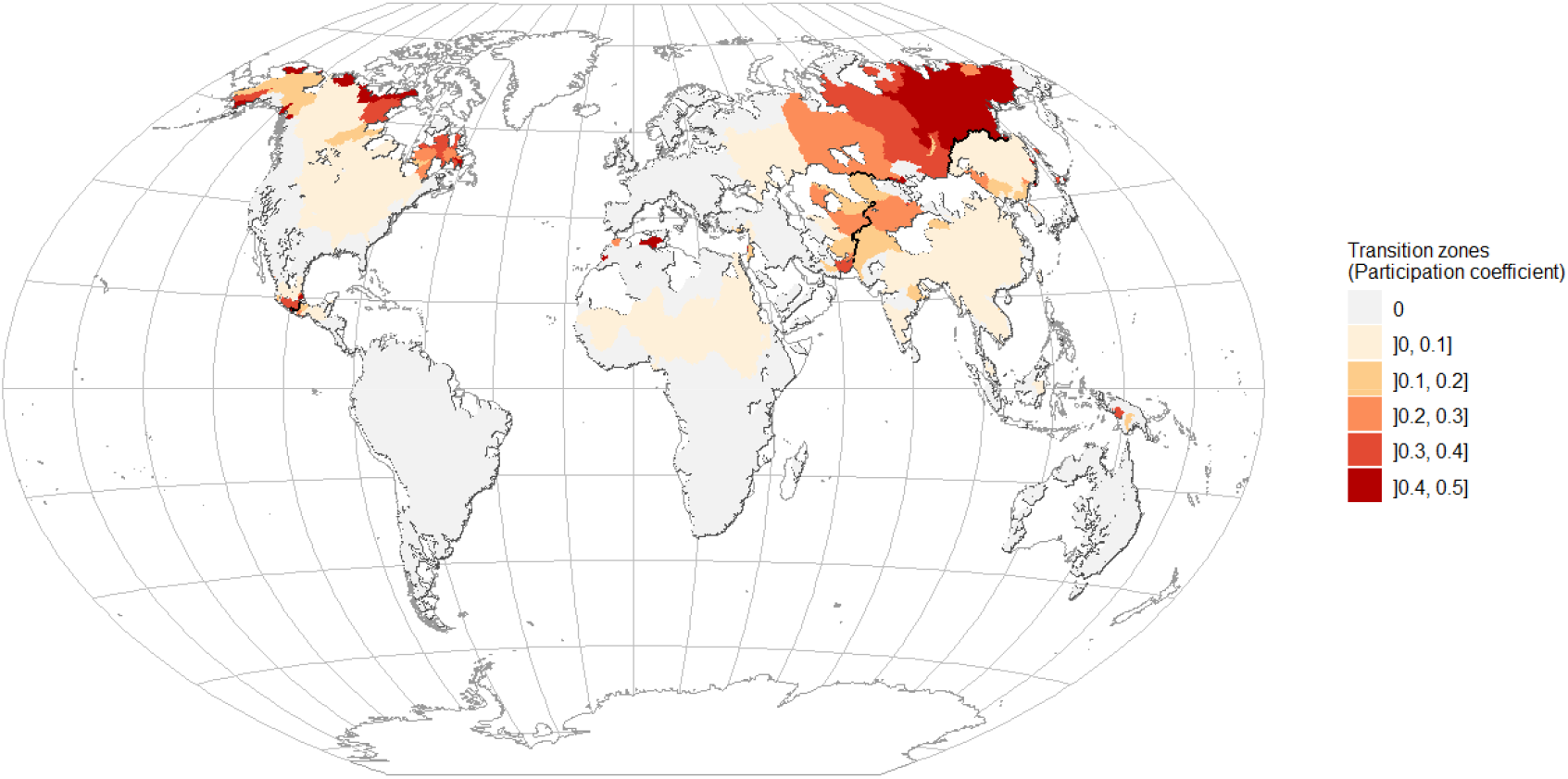
Map of transition zones between regions (level 2) characterised by the participation coefficient of basins, *i.e* the proportion of species in a basin that come from regions other than the region of this basin. Thick black lines indicate frontiers, which are shared between two neighbouring regions.

Overall, the species that were distributed across boundaries could be separated in two broad categories (Appendix S4). First, we found that the majority of shared species had restricted distributions close to regional boundaries, with occasional occurrences beyond. For example, the two-spot livebearer *Heterandria bimaculata* has a distribution endemic to Central America at the northernmost part of the Neotropical region and incurred in two basins of the Nearctic. Second, we found a limited number of species with large spatial distributions that were able to incur in multiple basins of other regions. The best example is the Northern pike *Esox lucius* which is one of the two species distributed across both supercontinental regions. Another example is the Eurasian minnow *Phoxinus phoxinus* that is widespread in the Palearctic with multiple occurrences in the Sino-Oriental region.

On the other hand, we found almost no transition zones at the boundaries of the Ethiopian or the Australian regions. For the Ethiopian region, only a few basins in northern Africa and the Middle East shared species between the Ethiopian and Palearctic regions (Figure 4) as illustrated by the limited number of links on the network (Figure 3). Only nine species were shared between these two regions (Appendix S4). For the Australian region, only three species dispersed through the Wallace-Huxley boundaries (Figure 3, Appendices S4 and S6): two species assigned to the Oriental region (*Aplocheilus panchax* and *Datnioides polota*) occur in Papuasia, and one species assigned to the Australian region (*Ophisternon bengalense*) was distributed in Australia, Papuasia and Southeastern Asia.

## Discussion

Here, we provide the first global bioregionalisation of freshwater fishes based on a quantitative analysis of species distributions. We found that the freshwater fish world is first divided into two supercontinental regions, the Old World and New World. Nested within these two supercontinental regions we found six major biogeographical regions with very high degrees of endemism: the Old World was divided in the Sino-Oriental, Ethiopian, Palearctic and Australian regions, whereas the New World was composed of the Neotropical and Nearctic regions. Nested within each of these major biogeographical regions we found subregions with varying degrees of endemism. Some regions had a few large subregions, while others had numerous subregions of heterogeneous size, species diversity and endemism. We found transition zones between the Nearctic and Neotropical regions, and between the Palearctic and Sino-Oriental regions. Furthermore, we found the Ethiopian and Australian regions to have almost no transition zones with other regions. The fish species that were distributed across regions were of two types: (i) species with ranges that were restricted to the proximity of the regional boundaries with occasional overlaps across boundaries and (ii) species with large distributions that expanded their range beyond boundaries.

Our results compellingly contradict our initial hypothesis that freshwater fish may have biogeographical regions different from the terrestrial vertebrate scheme of Sclater-Wallace because of their restricted dispersal abilities and the specific spatial-temporal dynamics of riverscapes. Indeed, we found a total of six major biogeographical regions that were similar in size and location to the six biogeographical regions identified by Wallace (1876). The only differences were the locations of several boundaries. More surprisingly, our regions were also similar to the coarse biogeographical regions based only on freshwater fish families identified by Matthews (1998) and that also included diadromous fish species. Our Neotropical region is the only major difference from results obtained by Matthews (1998) who identified a distinct cluster south to the Andes. This difference may be explained by the fact that this area was mostly colonised by the family of Galaxiidae that migrates between freshwater and oceans, and consequently have not been included in our analysis.

In addition, we also identified transition zones broadly following the bioregionalisation model of Morrone (2015), suggesting that freshwater fish regions were shaped by vicariance and geodispersal events similar to other groups. However, we observed extremely high rates of endemism for each region, more than 96% of endemic species for all regions except the Palearctic (89%). These endemism rates far exceed the endemism rates for other continental vertebrates such as birds (11 to 84% endemism in major regions, see Appendix S8), mammals (31% to 90%), herptiles (amphibian and reptiles: 46 to 95%; amphibian only: 66 to 98%), as calculated for regions of Procheş & Ramdhani (2012, Table S8.1 in Appendix S8) and Holt et al. (2013, Table S8.2 in Appendix S8). Freshwater fish endemism rates also exceed rates reported in marine biogeographical realms (17 to 84%, Costello et al., 2017). Therefore, we conclude here that freshwater fishes are likely to have among the highest rates of species endemism for major biogeographical regions.

The two major supercontinental regions we identified might seem to contradict Morrone’s biogeographical kingdoms (2015). Indeed, Morrone (2015) hypothesised that three major kingdoms could be derived from formerly disconnected land masses (i.e., Holarctic, Holotropical and Austral). This apparent contradiction is explained by our analyses at the species level which was not designed to reflect ancient biogeographical kingdoms resulting from the Gondwana split. Rather, the two supercontinental regions described here suggest that (1) the six major biogeographical regions have merged into two super-continents with biotic exchanges and (2) that strictly freshwater faunas have been isolated between these two super-continents for the past 10 to 20 million years. Only two species are shared between supercontinental regions, and genetic studies indicate that the only species documented, the pike *Esox lucius*, likely expanded its range after recent glaciations (Maes, van Houdt, De Charleroy, & Vockaert, 2003; Jacobsen, Hansen, & Loeschcke, 2005). This current pattern of isolation of the two supercontinental regions will have far-reaching implications for the future evolution of freshwater fish faunas. The imprint of this isolation on fish faunas will be visible for millions of years, likewise to the historical imprint of the split of Pangea that can be seen today on phylogenetically-informed bioregionalisations (or bioregionalisations at a higher taxonomic level such as families; e.g. Morrone et al. 2015). This pattern is, as far as we know, specific to strictly freshwater fishes. We can expect a different pattern for diadromous fishes which can disperse through marine waters and thus occur in both the Nearctic and the Palearctic (e.g., species in genera *Salmo, Salvelinus, Cottus, Lotta, Pungitius, Gasterosteus*).

Therefore, our hierarchy of supercontinental regions, regions and subregions should not be seen as the hierarchy of freshwater fish diversification history (i.e., our species-level supercontinental regions do not bear the same meaning as kingdoms in Morrone et al. 2015), but rather as the *recent* biogeography of strictly freshwater fish faunas. This recent biogeography illustrates spatially-nested supercontinental regions, regions and subregions all characterised by very high degrees of endemism. These very high degrees of endemism reflect the strong isolation of strictly freshwater fish faunas because of the lack of dispersal during the Neogene (caused by the isolation of the different land masses during the late Cretaceous to Paleogene) and the limited magnitude of fish dispersal events that occurred after land masses merged during the late Miocene to early Pleistocene. Only 86 (0.76%) of the world’s 11 295 species can be found across supercontinental or regional boundaries Therefore, 99.24% of the world’s freshwater fishes occur in a single supercontinental region and a single region, which ascertains the biological reality of our delineated clusters.

### Regions and transition zones

The Neotropical and Nearctic regions of freshwater fishes are very similar to the Neotropical and Nearctic regions highlighted for terrestrial mammals, amphibians and bird species (Kreft & Jetz, 2010; Procheş & Ramdhani, 2012; Holt et al., 2013). These two regions are therefore in agreement with the synthetic biogeographical regionalisation of Morrone (2015). As for other groups of organisms, we identified a Mexican transition zone between Nearctic and Neotropical regions, suggesting that these organisms were affected by biotic interchange between the Americas, but to a much lesser extent than terrestrial vertebrates (Bussing, 1985; Smith & Bermingham, 2005). Indeed, we found that only 25 species with restricted distributions were shared between Nearctic and Neotropical regions, as illustrated by the restricted area of the transition zone (Figure 4). These species belong to both primary and secondary freshwater families, which colonised Mesoamerica separately, as suggested by molecular analyses (Smith & Bermingham, 2005). Secondary freshwater fishes probably dispersed through Mesoamerica before the formation of the Panama isthmus, during periods of high runoff leading to temporary freshwater or brackish-water bridges in marine waters coupled with northward discharge of the proto-Amazon during the Miocene ∼ 18-15 Ma (Smith & Bermingham, 2005; Hoorn et al., 2010). Later, primary freshwater families dispersed through Mesoamerica during the Isthmus formation via landscape diffusion (Smith & Bermingham, 2005). Subsequent changes in the landscape led to increasing isolation of basins within Mesoamerica as well as occasional connectivity events, which in turn shaped the regional patterns of dispersal and diversification (Smith & Bermingham, 2005; Dias et al., 2014) that probably drove the restricted extent of the Mexican transition zone for freshwater fishes.

The Ethiopian region of freshwater fish resembles the Ethiopian regions of terrestrial vertebrates except for its northern limit. We found that this region expanded beyond the limits of the Sahara up to the Mediterranean sea, similarly to flightless terrestrial mammals (Kreft & Jetz, 2010). However, flightless terrestrial mammals were also found to expand their Ethiopian boundary beyond the Arabian Peninsula into Central Asia, which was not the case here. Moreover, most studies for terrestrial vertebrates located the northern boundary of the Ethiopian region south of the Sahara (Procheş & Ramdhani, 2012; Rueda et al., 2013; Holt et al., 2013; Vilhena & Antonelli, 2015). This discrepancy may be explained by several factors. First, most studies on vertebrates identified the Sahara and the northern coast of Africa as transition zones between the Palearctic and Ethiopian regions (Holt et al., 2013; Morrone, 2015). This transition zone was not identified in our analyses: Palearctic fishes only anecdotally crossed the Mediterranean Sea, while their African counterparts merely ventured into the Middle-East. Therefore, we can hypothesise that the Mediterranean Sea was an insurmountable barrier for freshwater fishes, contrary to terrestrial vertebrates that could disperse through Straits of Gibraltar. Dispersal during the desiccation of the Mediterranean Sea during the Messinian salinity crisis about 6 Ma has been proposed with the Lago Mare hypothesis (Bianco, 1990). This hypothesis stated that, during the refilling of the Mediterranean Sea, a freshwater or brackish phase occurred which would have permitted large scale dispersal of freshwater fishes across the Mediterranean basin. However this hypothesis received limited support in the light of phylogenetic studies (Levy, Doadrio, & Almada, 2009; Perea et al., 2010). Hence, freshwater fish dispersal during the Messinian salinity was not the same as that of other groups (e.g., Veith *et al*. 2004; Agustí *et al*. 2006). A probable dispersal pathway for freshwater fishes was through diffusion (*sensu* Lomolino et al., 2016) between the Nile and Mediterranean basins of the Middle East (illustrated by the distributions of *Clarias gariepinus* and four African cichlid species). This Middle-East dispersal pathway corroborates to some extent the Saharo-Arabian transition zone proposed by Morrone (2015). This dispersal pathway was identified for the early colonisation of Cyprinidae from the Sino-Oriental to Ethiopian region (Gaubert, Denys, & Oberdorff, 2009). Second, drainage basins north of the Sahara are extremely poor in freshwater fish species, notably because of their arid nature. Northernmost basins were probably colonised by Ethiopian species during repeated periods of aquatic connectivity across the Sahara region, the most recent being the Holocene African Humid period (∼11 to 8ka) (Drake, Blench, Armitage, Bristow, & White, 2011).

The Sino-Oriental region is a distinctive feature of freshwater fishes, for two reasons. Firstly, the northern boundary between Sino-Oriental and Palearctic is located beyond the Himalayas and the Gobi desert to the North-West, and goes up to the Stanovoï mountains located north of China, whereas all other groups are limited to a boundary extending longitudinally from the Himalayas to the China Sea (Cox, 2001; Procheş & Ramdhani, 2012; Rueda et al., 2013; Holt et al., 2013; Morrone, 2015). Secondly, this region is composed of two major subregions that were frequently identified as distinct regions in our sensitivity analysis. In particular, the Sinean subregion hosting a rich freshwater fish fauna (963 species) with 72% of endemics seems unique to freshwater fishes, and has been identified more as a transition zone for other groups (Procheş & Ramdhani, 2012; Holt et al., 2013; Morrone, 2015). We can speculate that this characteristic Sino-Oriental region for fishes arose from several factors. Firstly, the entire region was prone to fish speciation since the Eocene (55My), probably because of the very high diversity of aquatic habitats combined with the repeated rearrangement of rivers through capture and glaciation melting (Dias et al., 2014; Kang et al., 2014; Kang, Huang, & Wu, 2017; Xing, Zhang, Fan, & Zhao, 2016). For example, the Cyprinidae family, which accounts for 45% of Sino-Oriental species, originated from the Indo-Malaysian tropical region and has likely radiated into Asia since the Eocene (Gaubert et al., 2009). Secondly, the northern boundary is located farther North than from other groups, suggesting that mountain barriers were more important in defining boundaries for fishes than for other groups, whose boundaries appeared to be rather defined by a combination of tectonics and climate (Ficetola et al., 2017). Consequently, the fish transition zone between Sino-Oriental and Palearctic is not located near areas of recent tectonic merging as reported for other groups (Morrone, 2015). In addition, this transition zone is asymmetrically distributed towards the Palearctic (Figure 4) and possibly exceeding the asymmetry reported for other groups (Sanmartín, Enghoff, & Ronquist, 2001), probably because of the extreme differences in fish richness between both regions. Given the mountainous nature of the boundary, dispersal pathways probably emerged at river confluences when the sea level dropped (Dias et al., 2014), both at the Northeastern and Southwestern parts of the Sino-Oriental boundaries (Gaubert et al., 2009).

The Australian region is the most depauperate of all fish regions, probably owing to the combination of the complete isolation of this region for the last 60 million years (Scotese, 2016) and the dryness of the Australian continent. The boundary between Australian and Sino-Oriental encompasses the entire Wallacea: the Sino-Oriental extends to the limits of the Sunda shelf, whereas the Australian extends to the limits of the Sahul shelf (Lomolino et al., 2016). Only three species occur on both sides of the boundary. Two Sino-Oriental species (*Aplocheilus panchax* and *Datnioides polota*) are Sino-Oriental species known from one drainage basin of the Australian region, and one species (*Ophisternon bengalense*) is distributed in 10 and 11 basins of Australian and Sino-Oriental respectively. *O. bengalense* lives in estuaries and is tolerant to brackish waters, and thus may have dispersed through Wallacea (e.g., Capobianco & Friedman, 2018). On the other hand, the distributions of the other two species are more surprising, and the absence of data in all islands of Wallacea makes hypotheses highly speculative. These apparent disjoint distributions could be linked to unknown exceptional dispersal, undocumented species introductions, or misidentification (i.e. two species misidentified as a single species).

### Subregions

At subregional and finer levels, we found multiple clusters with varying degrees of endemism, with a substantial number of species distributed between clusters (see Figure 3). While it indicates that strictly freshwater fish species display strong endemism patterns at subregional spatial scales, it also suggests a reticulated history of river basins. The differences in number and endemism of subregions among major regions may be explained by the combination of habitat size and diversity, past climate change, and paleoconnectivity during the Last Glacial Maximum (LGM, see Leprieur *et al*. 2011; Tedesco *et al*. 2012; Dias *et al*. 2014). Past climate change had an enormous impact on high-latitude regions, such as the Nearctic and northern parts of the Palearctic (Leprieur et al., 2011). Most of these areas were covered by ice sheets during the LGM. These Northern areas were colonised by species after the LGM (e.g., Rempel & Smith 1998) from refuges located in the southernmost parts of these regions (Mississipi basin for the Nearctic, Danube basin for the Palearctic). Consequently, the relatively recent re-colonisation explains these large species-poor subregions. On the other hand, such climate events were less extreme in tropical regions thereby allowing lineages to thrive for a long period (Tedesco, Oberdorff, Lasso, Zapata, & Hugueny, 2005). The combination of this prosperity with the high diversity and size of habitats in tropical regions, the long-term isolation of drainage basins during the Pleistocene (Dias et al., 2014) as well as stable climatic history and favourable climatic conditions (Wright, Ross, Keeling, McBride, & Gillman, 2011) probably generated conditions favourable to divergence and radiation processes in tropical subregions. This last hypothesis may explain the numerous tropical subregions with high diversity and endemism we found. The only apparent contradiction could arise from the two large subregions with high endemism of the Sino-Oriental. However, these two subregions probably reflect the uplift of the Tibetan plateau that led to their isolation (Kang et al., 2014). In turn, these two subregions included numerous smaller ones with high diversity and endemism (see Figure S7.15) similar to the other tropical regions.

### Robustness of findings

This first global quantitative analysis of the biogeography of freshwater fishes is based on a large scale database compiling occurrence data from thousands of sources and is thus inevitably subject to errors and incomplete data (Tedesco et al., 2017). To minimise errors, a careful screening and correction procedure has been implemented for this database (see Tedesco et al., 2017). Reassuringly, the results obtained from other clustering methods as well as our sensitivity analysis suggested that the regions we identified are robust. Furthermore, regions were all spatially coherent (even though no spatial information was provided at any stage of the process) for the first two levels, with high degrees of endemism, indicating the quality of both the dataset and the bioregionalisation approach.

The network method assigns clusters to species (as explained in the methods), which is an asset over distance-based clusters. However, one major caveat needs to be acknowledged. Species are assigned to the region where their present-day distribution is largest – this region is not necessarily the region where they originated from. A perfect example is the Characidae family at the transition between Nearctic and Neotropical: *Astyanax mexicanus* was assigned to the Nearctic, although its lineage is assumed to have colonised Mesoamerica during the Panama Isthmus formation (Smith & Bermingham, 2005).

Our results at the subregional scales have several limits. Firstly, while species introductions are relatively well documented between major continents or regions, we can expect that some human-assisted translocations of species among basins have not been documented at smaller spatial scales, thereby blurring subregional patterns of endemism. Secondly, heterogeneity in land topology led to vast differences in size and number of drainage basins within different geographical areas of the world. Furthermore, drainage basins (especially large ones) may have a reticulated history challenging their validity as biogeographical units at sub-regional levels (e.g., see Dagosta & Pinna, 2017 and references therein). Third, for areas with numerous small basins, species lists were not necessarily available for all of them, and thus identification of provinces beyond the sub-regional scale would be speculative. Likewise, fine-scale data was not available in similar quantity or quality within different regions of the world (e.g., remote areas of Africa or Papuasia remain poorly sampled compared to other regions). Last, the Map Equation is expected to identify transition zones as distinct clusters (Vilhena & Antonelli, 2015; Bloomfield et al., 2017). We did not observe this pattern at large scales, except for a few basins at the transitions between Nearctic and Neotropical regions or between Sino-Oriental and Palearctic or Australian regions. However, at sub-regional and finer scales, transitions are likely to stand out as separate zones (Bloomfield et al., 2017), which may not necessarily be an appealing property since the participation coefficient is informative enough to describe transition zones. For all these reasons, we deemed preferable not to investigate our results below the third level, as such fine-scale provinces would be better studied in regional studies (e.g., Smith & Bermingham, 2005; Kang et al., 2014).

### Concluding remarks

This first quantitative study of freshwater fish bioregions revealed that their biogeography was probably shaped by the same major drivers as other continental groups of organisms, with peculiar exceptions such as the Sino-Oriental region. These regions identified with species distributions probably reflect relatively recent processes of dispersal and isolation. Ancient processes will be explored in future studies thanks to the newly available dated phylogenies of actinopterygian fishes (Rabosky et al., 2018).

We found that freshwater fishes, in addition to being the most diverse group of continental vertebrates, have extremely high rates of endemism, above 96% for all regions except the Palearctic. Furthermore, we found that tropical regions have a myriad of subregions with very high endemism and richness. These figures compellingly bespeak that freshwater fishes ought to be considered in hotspot analyses and raise many questions about the biogeographical consequences of the current high rates of freshwater fish introductions and extirpations (Villéger, Blanchet, Beauchard, Oberdorff, & Brosse, 2015).

## Acknowledgements

We thank Céline Bellard and Philippe Keith for useful discussions, and Aldyth Nyth and Lissette Victorero for English editing. We thank François-Henri Dupuich from derniercri.io for figure editing. Laboratoire Evolution et Diversité Biologique is part of the French Laboratory of Excellence projects “LABEX TULIP” and “LABEX CEBA” (ANR-10-LABX-41, ANR-10-LABX-25-01).

## Biosketch

Boris Leroy is lecturer at the Muséum National d’Histoire Naturelle of Paris. He is interested in the biogeography of aquatic organisms and how it is or will be impacted by global changes such as climate change and invasive alien species.

