## Appendix S2 for "Global biogeographical regions of freshwater fish species"

*Preprint manuscript submitted to the Journal of Biogeography*

**Appendix S2. Bioregionalisation results based on additional clustering methods**

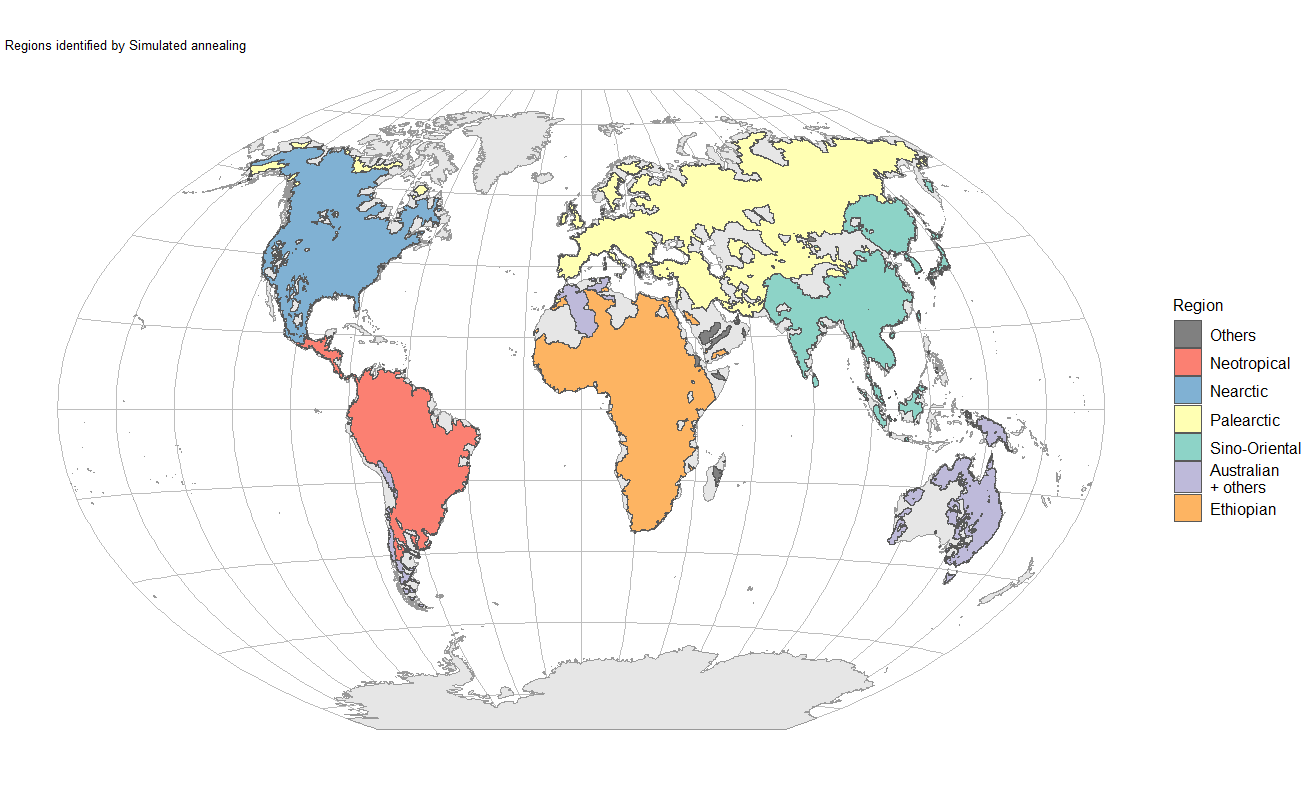
 **Figure S2.1** Biogeographical regions of freshwater fishes defined at the species level on the basis of the Simulated Annealing algorithm.

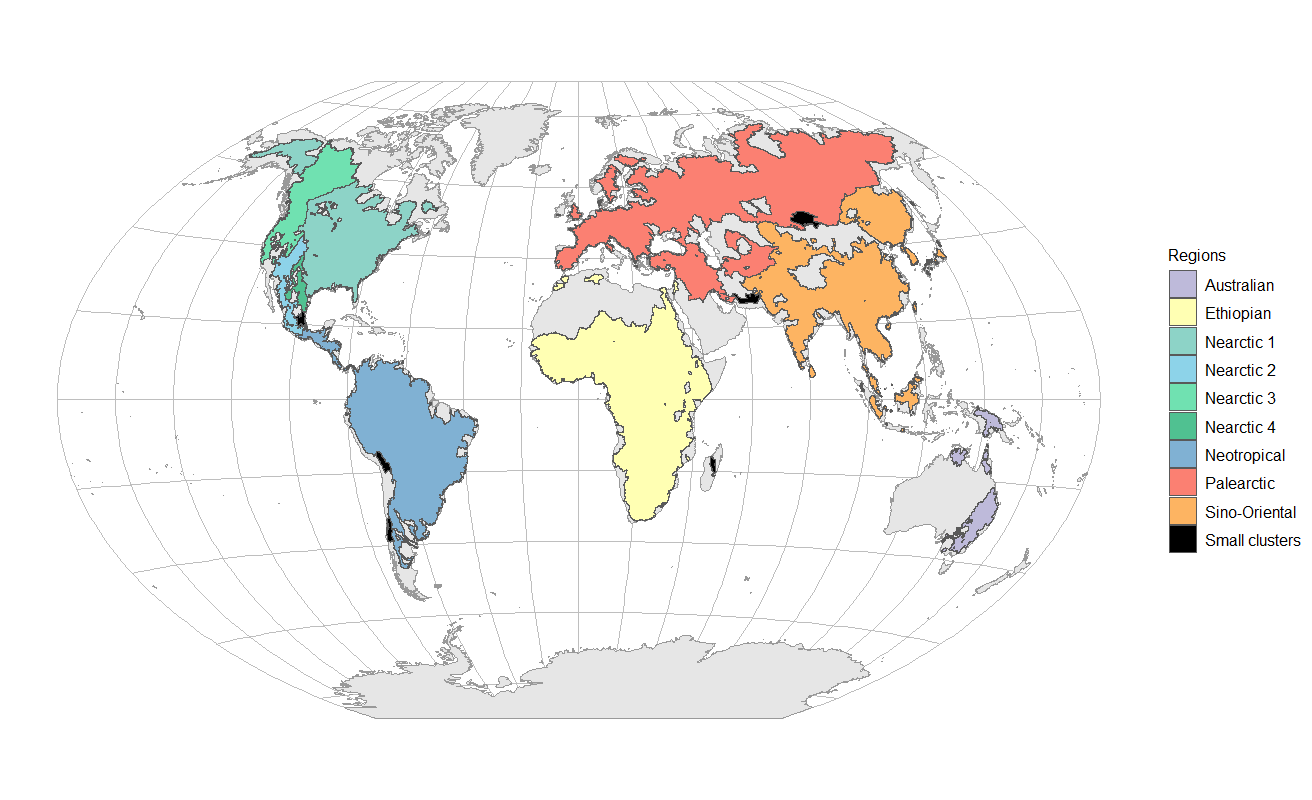

**Figure S2.2** Biogeographical regions of freshwater fishes defined at the species level on the basis of β_sim_ distances between basins and UPGMA classification. Regions were identified as clusters with a 0.998 cut-off. Drainage basins with less or equal than five species were removed for this analysis.

**Methods and results for the Simulated Annealing regionalisation**

We applied the Simulated Annealing method as described in Bloomfield et al. (2017) on the exact same network as for the Map Equation method. We used the netcarto function of the rnetcarto R package version 0.2.4 (Doulcier & Stouffer, 2015) with default settings.

The results from the Simulated Annealing algorithm were very similar to the results obtained from the hierarchical Map Equation algorithm at level 2. Two differences were observed: first, the Central Asian portions of the Sino-Oriental were assigned to the Palearctic. This area also corresponds to the area that was occasionally defined as a transition zone in our sensitivity analysis (see Appendix S3). Therefore, this pattern is likely due to the fact that the area is a relatively species poor transition zone with species from both regions. Second, some of the most species poor basins of the Ethiopian and Neotropical zone are associated with the Australian zone, even though they do not have any species in common. This aberration is likely an artefact of the method.

**Methods and results for the β diversity regionalisation**

We followed the framework described in Kreft & Jetz (2010) to define biogeographical regions on the basis of a measure of species turnover. We chose the turnover component of the Sorensen β diversity index (β_sim_) following Leprieur & Oikonomou (2014) because this index is assumed to be less dependent on differences in species richness. We applied a hierarchical clustering algorithm on the compositional dissimilarity matrix: we used the Unweighted Pair-Group Method using arithmetic Average (UPGMA) as recommended by Kreft & Jetz (2010). However, as demonstrated in Dapporto et al., (2013), the order of rows in the dissimilarity matrix can heavily influence the UPGMA dendrogram, likewise to all other clustering algorithms proposed in Kreft & Jetz (2010). Note that this aspect has seldom been investigated in bioregionalisation studies despite its potentially important impact on the results. Our preliminary randomization procedures indeed revealed that changing the order of rows resulted in different dendrograms in terms of relations between regions. We therefore generated 1000 dendrograms by randomly shuffling the order of rows in the dissimilarity matrix, and we constructed a consensus tree with the recluster.cons function of the recluster R package (Dapporto et al., 2013). On the resulting consensus tree, we then explored the number of clusters that were obtained by cutting the tree at different threshold values ranging from 0.999 to 0.900 (Figure S2.3).

When all basins were included in the analysis, we obtained inconsistent results probably due to the numerous basins with low species richness, a well-known issue with hierarchical clustering (Kreft & Jetz, 2010). Indeed, we obtained three large clusters (cluster 1: South and Central America; cluster 2: North America, Europe, Asia and Africa; cluster 3: Australia) and several smaller clusters for cut-offs ranging from 0.999 to 0.925; and more than 1 000 small clusters for cut-offs below 0.925. Therefore, we removed all basins with richness below or equal to 5 and ran the clustering procedure again. With this species richness filter, we obtained eight large clusters and several small clusters for cutoffs ranging from 0.998 to 0.950 (Figure S2.2). The majority of the large clusters followed the clusters we obtained at level 2 (termed Regions) in our network analyses. The only major discrepancy was the Nearctic which was divided into four clusters. These four clusters could not be grouped together because all clusters were connected to the same node in the consensus tree (Figure S2.4), thereby preventing any hypothesis regarding biogeographical relations between regions with the β diversity method.

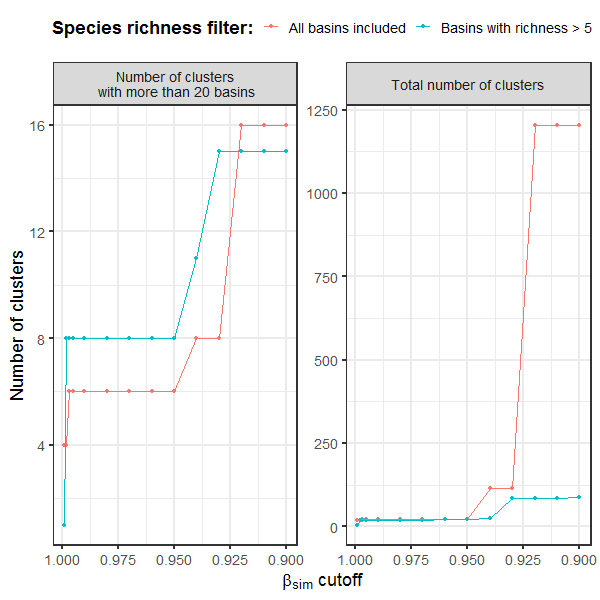

**Figure S2.3** Number of clusters obtained (y axis) by cutting the consensus tree at different cutoffs (x axis). The left panel shows the number of large clusters (number of basins ≥20) and the right panel shows the total number of clusters.

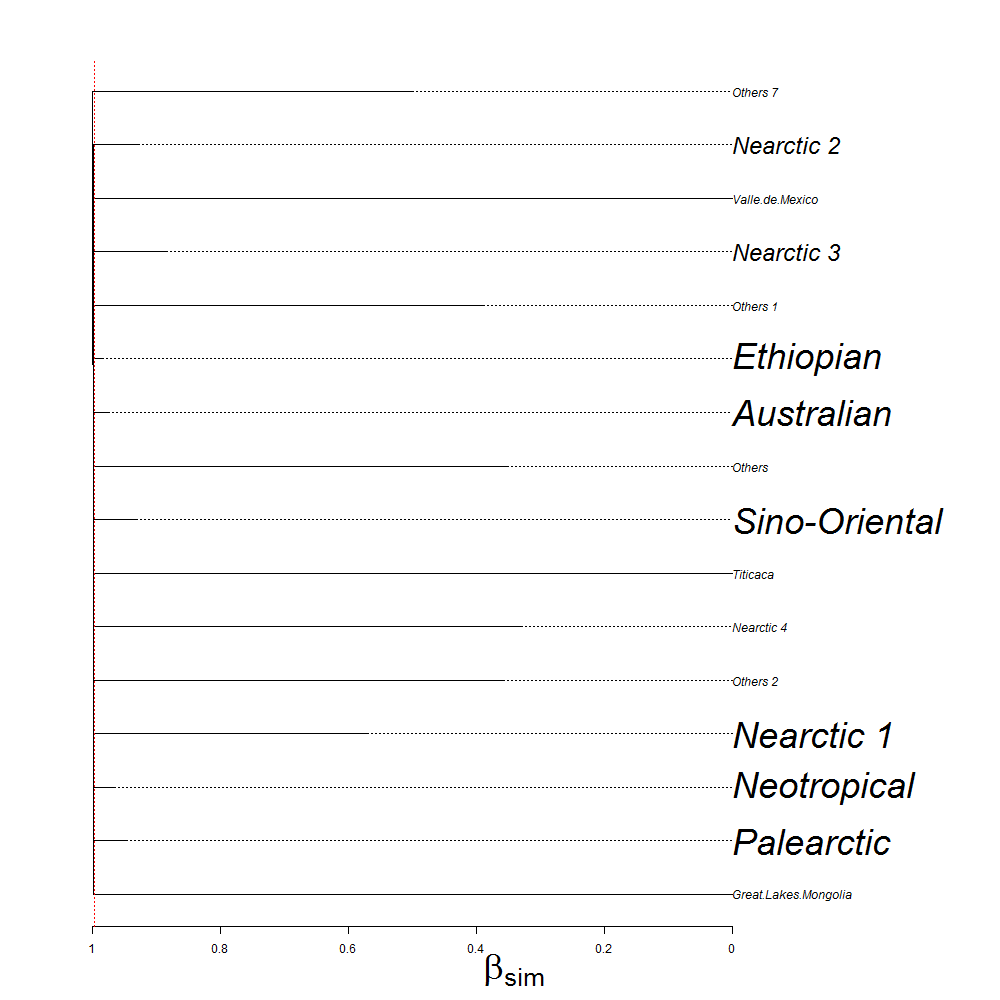

**Figure S2.4** Hierarchical consensus tree of freshwater fish basins. This hierarchical consensus tree was obtained by combining 1000 UPGMA trees on the dissimilarity matrix with rows randomly shuffled in each run. Because the tree is very large, we collapsed all nodes with values below 0.95 so that it is readable (the height of collapsed nodes is indicated by dotted lines). The number of basins for each collapsed node is indicated by the size of names: large names indicate nodes with more than 30 basins; intermediate names indicate nodes with 11 to 30 basins; small names indicate nodes with 1 to 11 basins. The vertical red dotted line indicate the cutoff (0.998) that we used to generate the map in Figure SX.2.
