## Appendix S3 for "Global biogeographical regions of freshwater fish species"

*Preprint manuscript submitted to the Journal of Biogeography*

**Appendix S3. Sensitivity analysis**

The Australian, Ethiopian, Nearctic, Neotropical and Palearctic regions were very robust because more than 95% of their original area was retrieved (Figure S3.5) in almost all simulations runs (see 10% quantile in Table S3.1). The Sino-Oriental region was the only region that was likely to significantly split, with two possible splits. The most likely split was the separation into two regions, the Sinean and Oriental (see box in Figure S3.5 and example map in Figure S3.6). A second, occasional split (5 runs out of 200) was the identification of the Central Asian part of the Sino-Oriental as a distinct region (see example map in Figure S3.7).

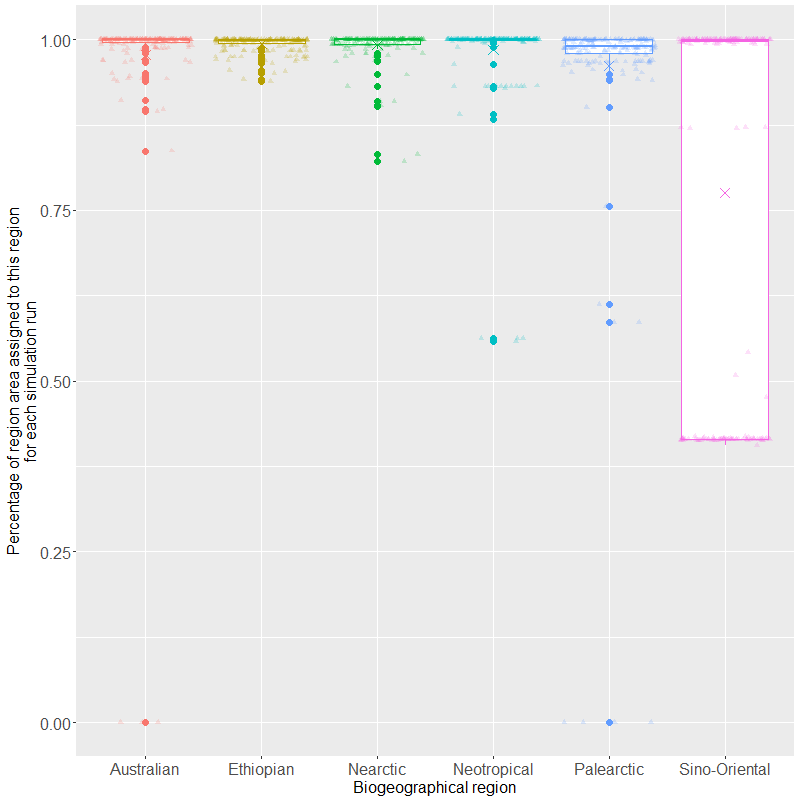

Sino-Oriental region split in Sinean and Oriental.

Figure S3.5. Box-and-whisker plots of the percentage of area of the original regions that were assigned to this same region in each of the 200 sensitivity analysis runs. Crosses represent average values; plain circles represent outliers and transparent triangles represent each individual run.

| **Table S3.1.** For the 200 sensitivity analysis runs, average and quantiles of the percentage of each region’s initial area that was retrieved. | | | | | | |
| --- | --- | --- | --- | --- | --- | --- |
|  | Sino-Oriental | Ethiopian | Palearctic | Australian | Neotropical | Nearctic |
| Average | 77.5% | 99.4% | 96.1% | 97.8% | 98.5% | 99.3% |
| Quantiles |  |  |  |  |  |  |
| 10.0% | 41.5% | 97.6% | 96.4% | 97.9% | 96.0% | 98.6% |
| 20.0% | 41.5% | 99.0% | 96.8% | 99.4% | 99.8% | 99.2% |
| 30.0% | 41.6% | 99.8% | 98.5% | 99.7% | 100.0% | 99.4% |
| 40.0% | 87.2% | 99.9% | 98.8% | 99.9% | 100.0% | 99.9% |
| 50.0% | 99.7% | 99.9% | 99.0% | 100.0% | 100.0% | 100.0% |
| 60.0% | 100.0% | 99.9% | 99.9% | 100.0% | 100.0% | 100.0% |
| 70.0% | 100.0% | 99.9% | 99.9% | 100.0% | 100.0% | 100.0% |
| 80.0% | 100.0% | 100.0% | 99.9% | 100.0% | 100.0% | 100.0% |
| 90.0% | 100.0% | 100.0% | 100.0% | 100.0% | 100.0% | 100.0% |

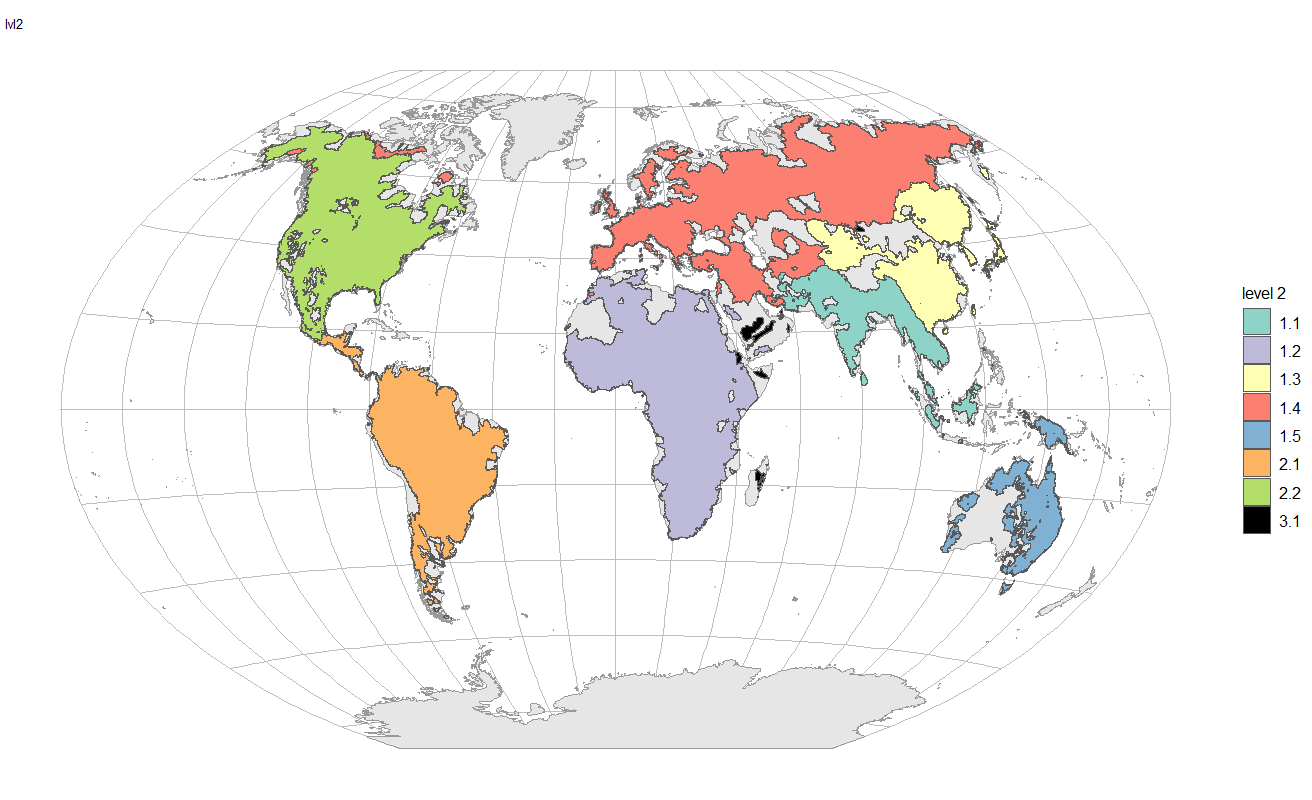

Figure S3.6. Example map of a simulation run (run #6) where the Sino-Oriental region was split into two regions, the Sinean region (region 1.3) and the Oriental region (region 1.1).

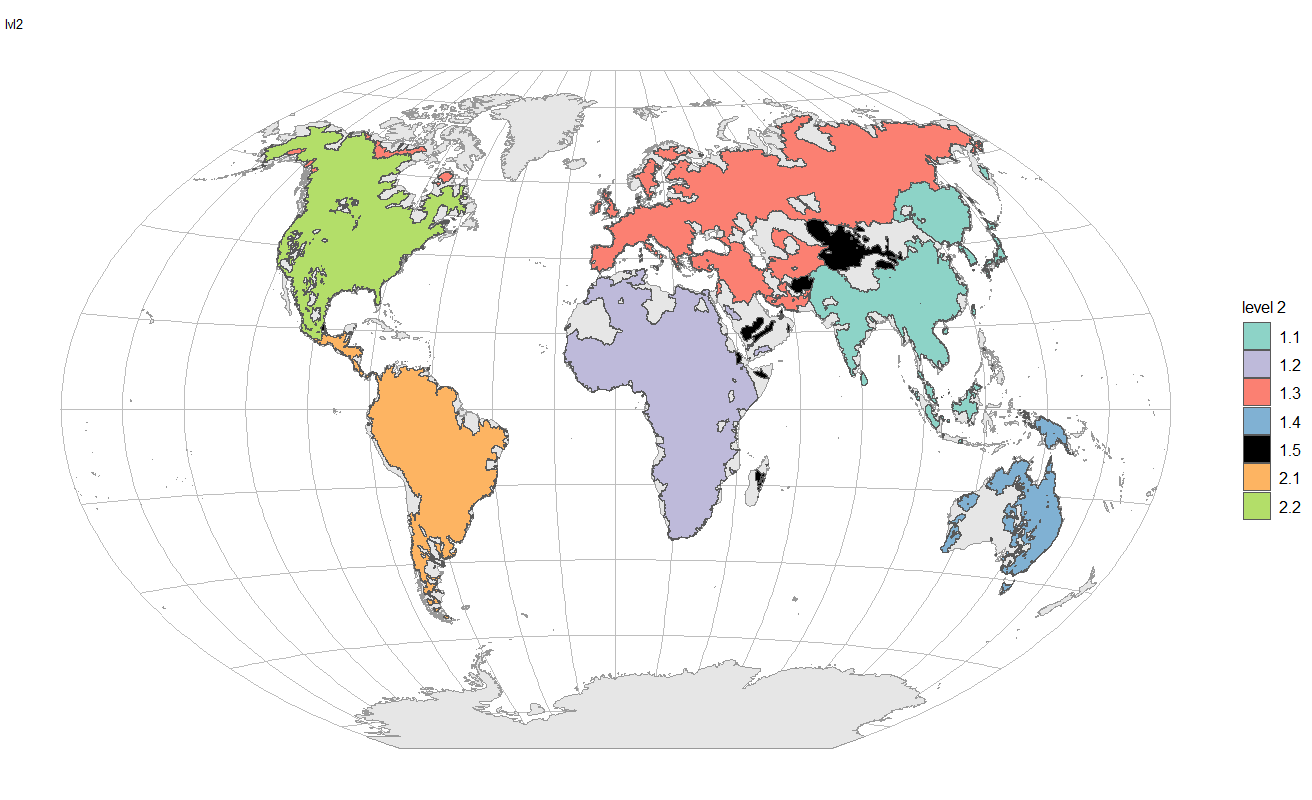

Figure S3.7. Example map of a simulation run (run #83) where ~13% of the Sino-Oriental region (mountainous parts of northwestern China, western Mongolia and east Central Asia) was identified as a distinct region.
