## Appendix S5 for "Global biogeographical regions of freshwater fish species"

*Preprint manuscript submitted to the Journal of Biogeography*

**Appendix S5. *Post-hoc* corrections to clusters.**

We corrected two types of inaccuracies in our bioregionalisation results. First, we found several basins of North America that were initially attributed to the Palearctic region (see northernmost parts of America in Figure S5.8a and b). These were composed of only one or two species, including the pike *Esox lucius*. Since *E. lucius* has been assigned to the Palearctic region, less-documented basins of North America where it is the only reported species (or with a single species of lower occurrence) were attributed to the Palearctic. Therefore, we manually reassigned all basins of North America to the New World at level 1, to the Nearctic at level 2, and to subregion “2.2.1”at level 3 (with the choice based on spatial coherence and species composition).

a. Supercontinental regions (level 1)

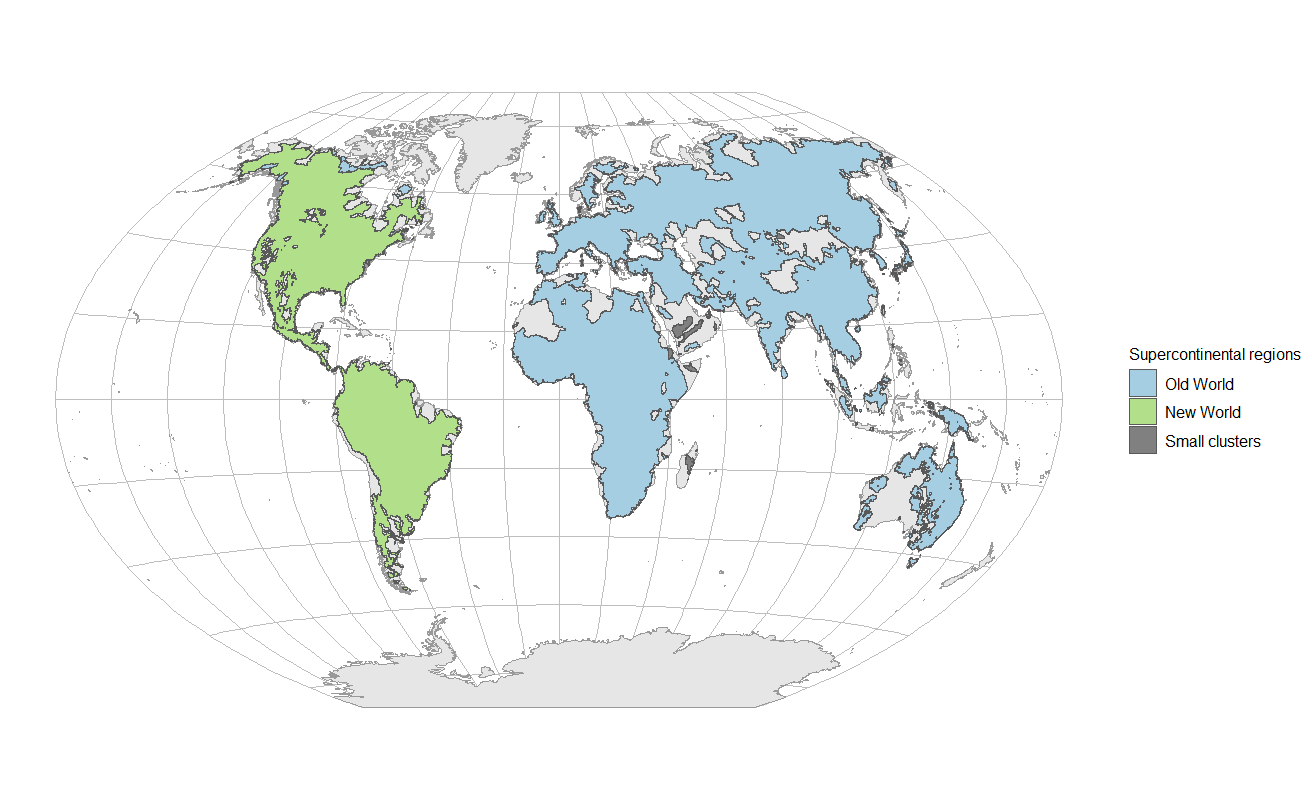

b. Regions (level 2)
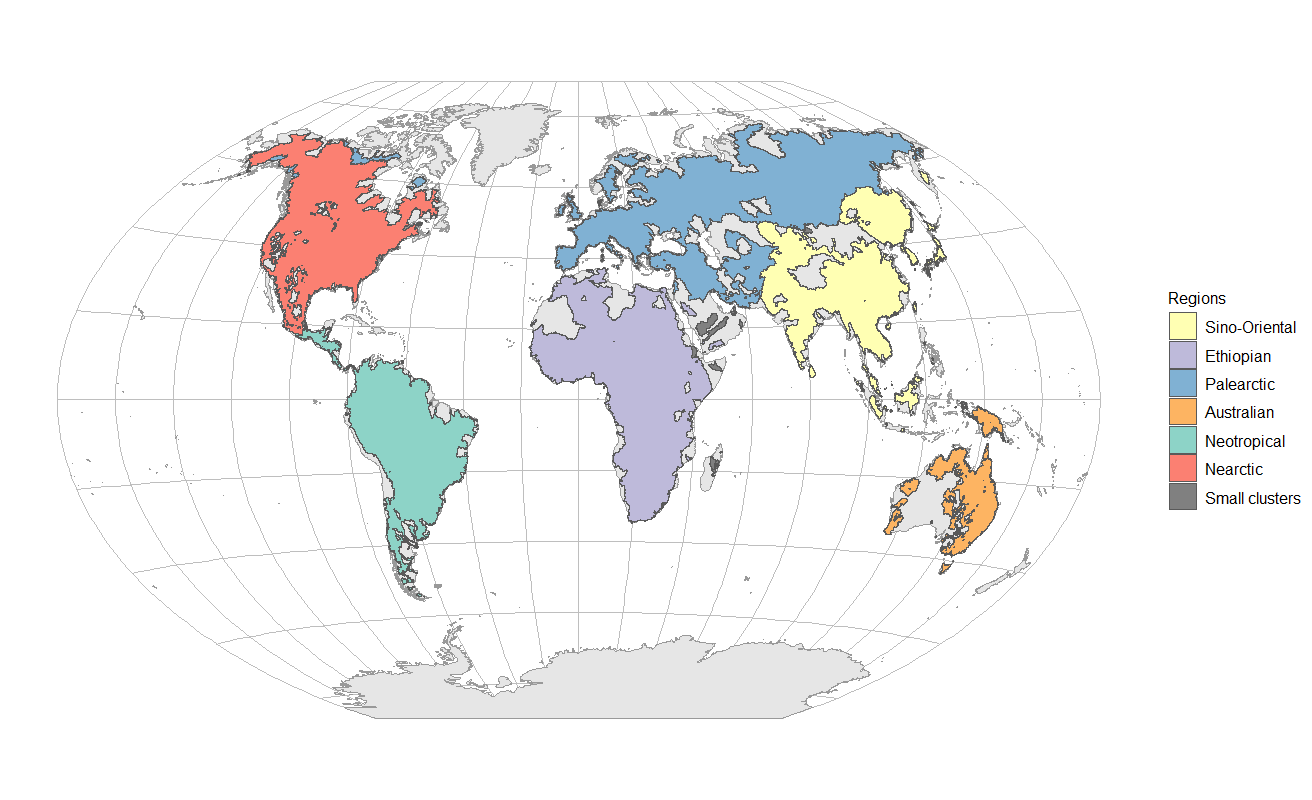

**Figure S5.8.** Uncorrected biogeographical regions of freshwater fishes defined at the species level with the Map Equation clustering algorithm.

Second, we found 14 tiny clusters at the first level (realm level), and an additional cluster at the second level (region level) which resulted in a total of 15 tiny clusters at level 2. Overall, these clusters represent 53 basins (0.7% of total surface of basins for which we have data), and 46 species (0.4% of total species richness of the database). We reviewed each of these clusters and manually corrected the hierarchy of regions. We provide below the rationale for their assignation. Cluster codes correspond to those provided by Map Equation. We provide a location map with species list for each cluster after the text.

- Cluster 3

This cluster exclusively included basins from Madagascar. All species and genera of this cluster are endemics, but one family (*Cichlidae*) is widespread throughout the Old World. Therefore, we decided to assign this cluster to the Old World (level 1) but kept it as a distinct, smaller region at level 2.

- Clusters 4 and 6

These two clusters are basins of the Arabian peninsula. They include a genus from the Palearctic (*Cyprinion*) but their species are endemic to the peninsula; they also include genera distributed in both the Ethiopian and the Palearctic (*Garra* and *Carasobarbus*). The balanced origins and distribution of these genera between the Ethiopian and the Palearctic lead us to consider that this region is probably an ancient transition zone. We decided to attribute these basins to the Palearctic, which, combined to the fact that other basins of the Arabian peninsula have been attributed to the Ethiopian, illustrates that the Arabian peninsula is probably a transition zone. Therefore, we assigned these cluster to the Old World at level 1, to the Palearctic at level 2, and to a subregion of their own at level 3.

- Clusters 5, 7, 10, 14, 15 and 16

These are Indonesian clusters on the Australian side with species of the genus *Melanotaenia* which is largely distributed in the Australian region, so we assigned these clusters to the Old World at level 1 and to the Australian region at level 2. At level 3, we assigned cluster 5 to subregion 1.4.2 and clusters 7, 10, 14, 15 and 16 to subregion 1.4.1 on the basis of geographical proximity.

- Cluster 8

This cluster is a unique basin with a unique endemic species in the Martinique island in the Caribbean. The genus *Anablepsoides* is distributed in the Neotropical so we assigned this cluster to the New World at level 1, Neotropical at level 2, and to a subregion of its own at level 3.

- Cluster 9

Cluster 9 is a complex zone in the African continent which was identified as a distinct biogeographical province (“Ethiopian Rift Valley province”) by Paugy (2010). We attributed this cluster to the Ethiopian region because Paugy (2010) considered that this is an “impoverished province of the Nilo-Sudan province”. Therefore, we assigned this cluster to the Old World at level 1, to the Ethiopian region at level 2, and to a subregion of its own at level 3.

- Cluster 11

This cluster is a Japanese cluster with a single endemic species whose genus *Rhodeus* is distributed in the Sino-Oriental. Therefore, we assigned this cluster to the Old World at level 1 and to the Sino-Oriental at level 2. At level 3, we assigned it to subregion 1.1.2 and at level 4 to subregion 1.1.2.1 on the basis of geographical coherence.

- Cluster 12

This cluster is the Gobi lake, at the boundary between the Palearctic and Sino-Oriental, with a single species of the genus *Barbatula*. The genus *Barbatula* is mostly Palearctic so we speculatively attributed this cluster to the Palearctic. Therefore, we assigned this cluster to the Old World at level 1, to the Palearctic region at level 2, and to the 1.3.23 subregion at level 3 on the basis of geographical coherence.

- Cluster 13

Cluster 13 is another cluster located on the African continent, with a single endemic species of the genus *Barbopsis*. Based on morphological analysis, Hayes & Armbruster (2017) suggest that this genus should be considered *Enteromius*, an African genus. Therefore, we assigned this cluster to the Old World at level 1, to the Ethiopian region at level 2, and to a subregion of its own at level 3.

Hayes, M. M., & Armbruster, J. W. (2017). The Taxonomy and Relationships of the African Small Barbs (Cypriniformes: Cyprinidae). *Copeia*, *105*(2), 348–362.

Paugy, D. (2010). The Ethiopian subregion fish fauna: An original patchwork with various origins. *Hydrobiologia*, *649*(1), 301–315.

1

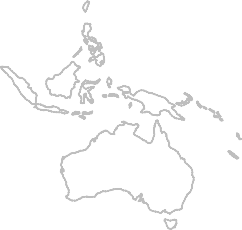

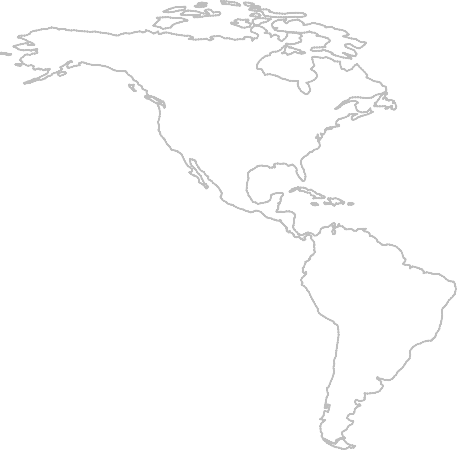

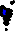

| **species.list** | **region** |
| --- | --- |
| Bedotia madagascariensis | 3 |
| Paratilapia polleni | 3 |
| Paretroplus kieneri | 3 |
| Paretroplus polyactis | 3 |
| Paretroplus tsimoly | 3 |
| Ptychochromis oligacanthus | 3 |
| Rheocles alaotrensis | 3 |
| Ancharius fuscus | 3 |
| Bedotia geayi | 3 |
| Bedotia marojejy | 3 |
| Rheocles pellegrini | 3 |
| Rheocles vatosoa | 3 |
| Gogo brevibarbis | 3 |
| Ptychochromis grandidieri | 3 |
| Katria katria | 3 |
| Oxylapia polli | 3 |
| Rheocles sikorae | 3 |
| Bedotia leucopteron | 3 |

2

3

4

5

6

7

8

9

10

11

12

13

14

15

16

17

18

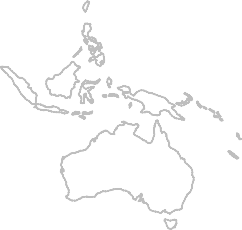

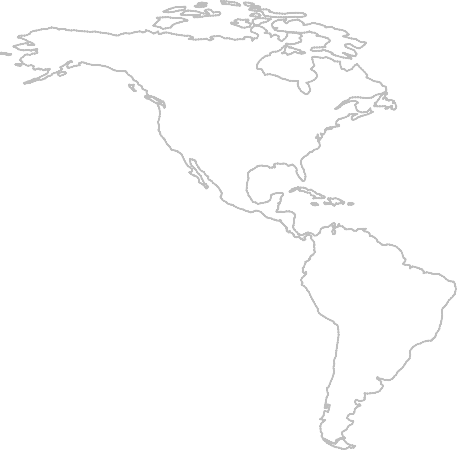

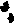

1

| **species.list** | **region** |
| --- | --- |
| Garra barreimiae | 4 |
| Cyprinion microphthalmum | 4 |
| Garra longipinnis | 4 |

2

3

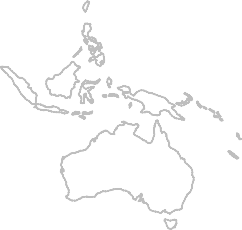

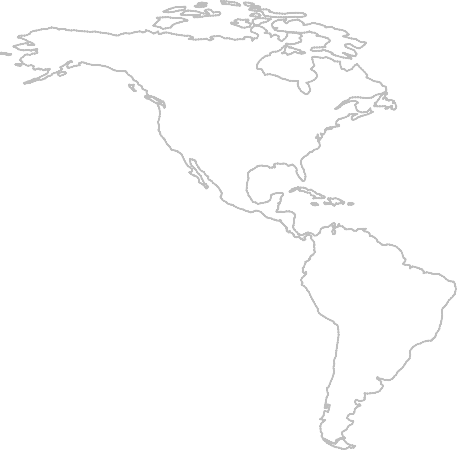

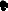

1

| **species.list** | **region** |
| --- | --- |
| Melanotaenia irianjaya | 5 |
| Melanotaenia ajamaruensis | 5 |
| Melanotaenia boesemani | 5 |
| Melanotaenia angfa | 5 |
| Melanotaenia parva | 5 |

2

3

4

5

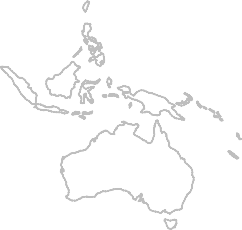

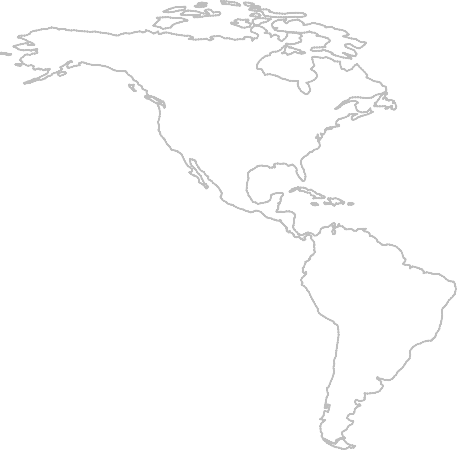

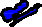

1

| **species.list** | **region** |
| --- | --- |
| Carasobarbus apoensis | 6 |
| Cyprinion mhalensis | 6 |
| Garra buettikeri | 6 |

2

3

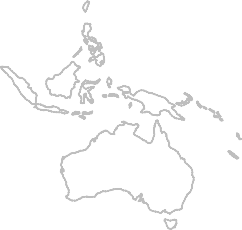

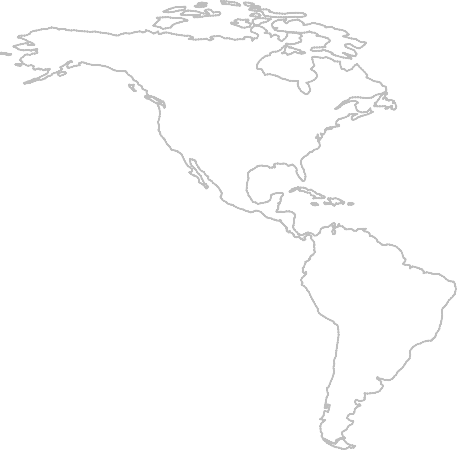

1

| **species.list** | **region** |
| --- | --- |
| Melanotaenia parkinsoni | 7 |

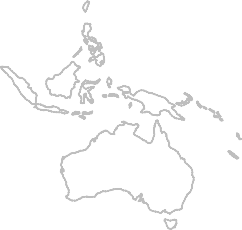

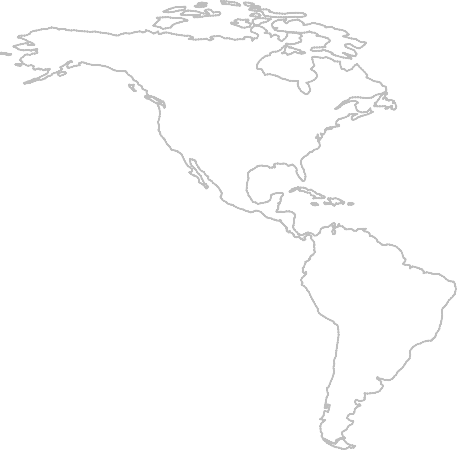

1

| **species.list** | **region** |
| --- | --- |
| Anablepsoides cryptocallus | 8 |

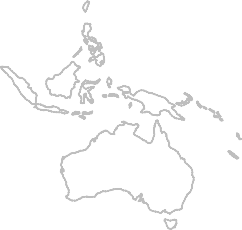

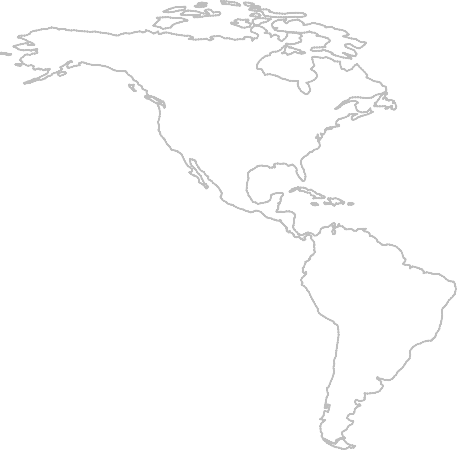

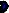

1

| **species.list** | **region** |
| --- | --- |
| Aphanius stiassnyae | 9 |
| Danakilia franchettii | 9 |

2

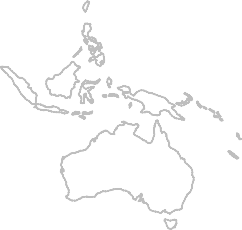

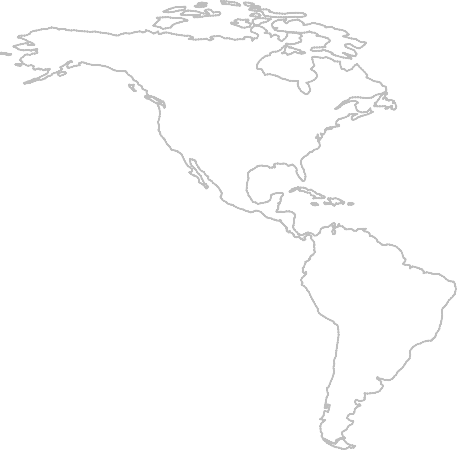

1

| **species.list** | **region** |
| --- | --- |
| Melanotaenia fredericki | 10 |

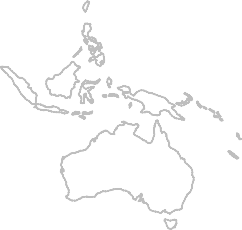

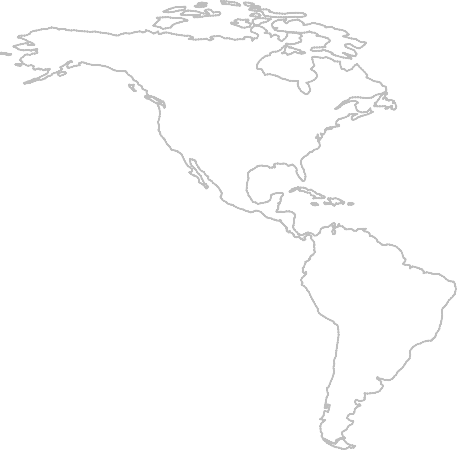

1

| **species.list** | **region** |
| --- | --- |
| Rhodeus suigensis | 11 |

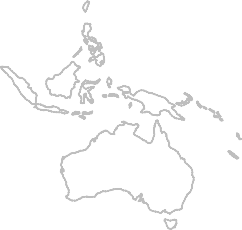

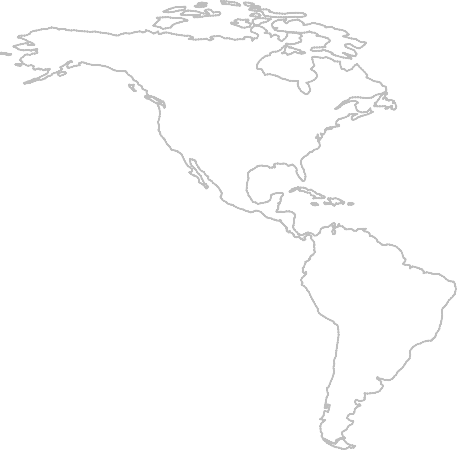

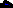

1

| **species.list** | **region** |
| --- | --- |
| Barbatula dgebuadzei | 12 |

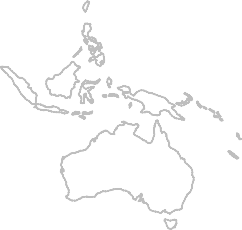

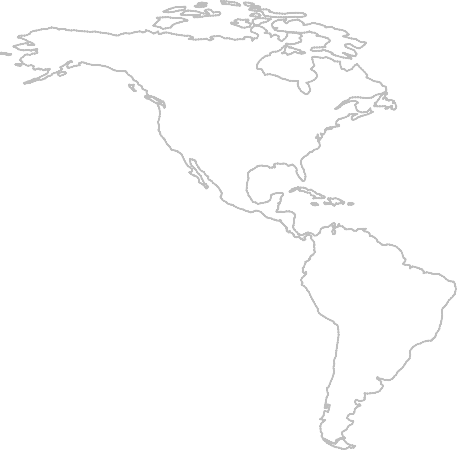

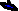

1

| **species.list** | **region** |
| --- | --- |
| Barbopsis devecchii | 13 |

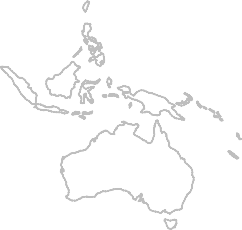

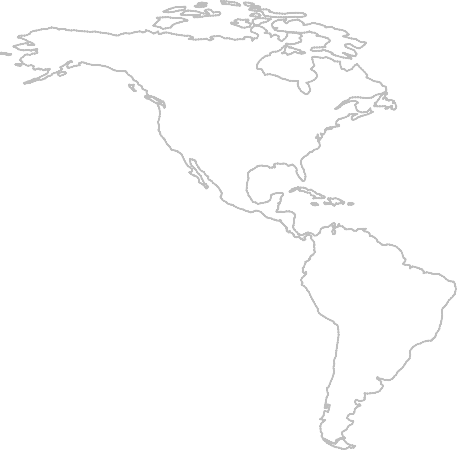

1

| **species.list** | **region** |
| --- | --- |
| Melanotaenia arfakensis | 14 |

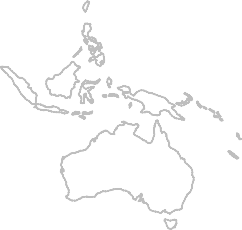

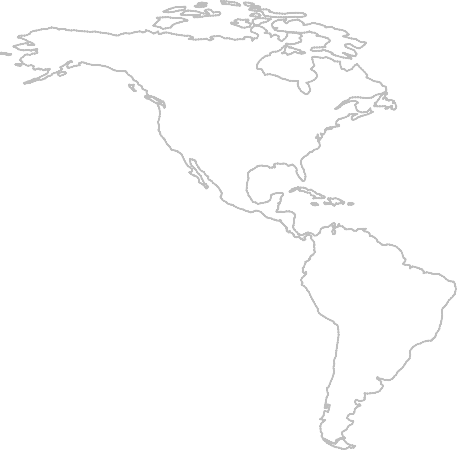

1

| **species.list** | **region** |
| --- | --- |
| Melanotaenia catherinae | 15 |

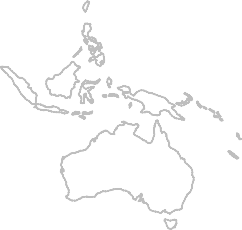

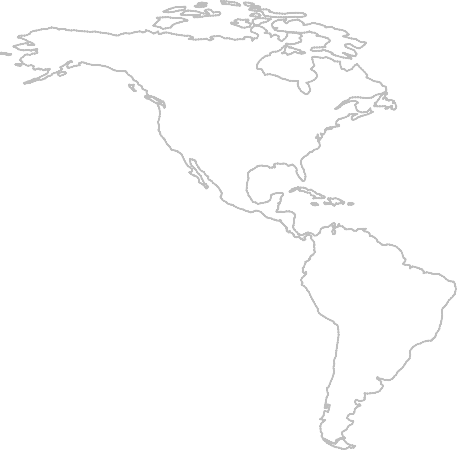

1

| **species.list** | **region** |
| --- | --- |
| Melanotaenia misoolensis | 16 |
