## Supplementary figures and images for "Global biogeographical regions of freshwater fish species"

### Appendix S6

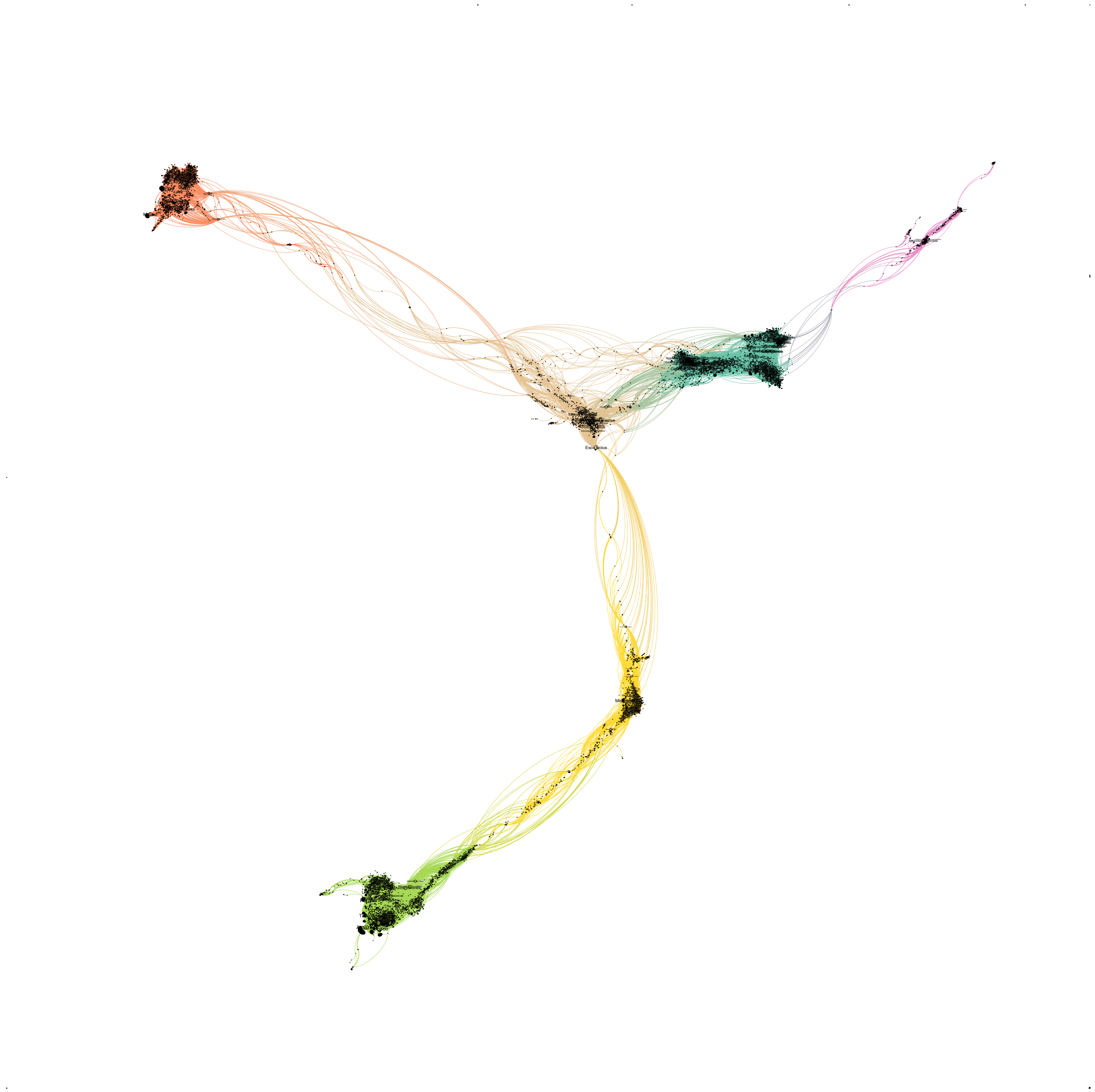
