## Appendix S7 for "Global biogeographical regions of freshwater fish species"

*Preprint manuscript submitted to the Journal of Biogeography*

**Appendix S7. Subregions**

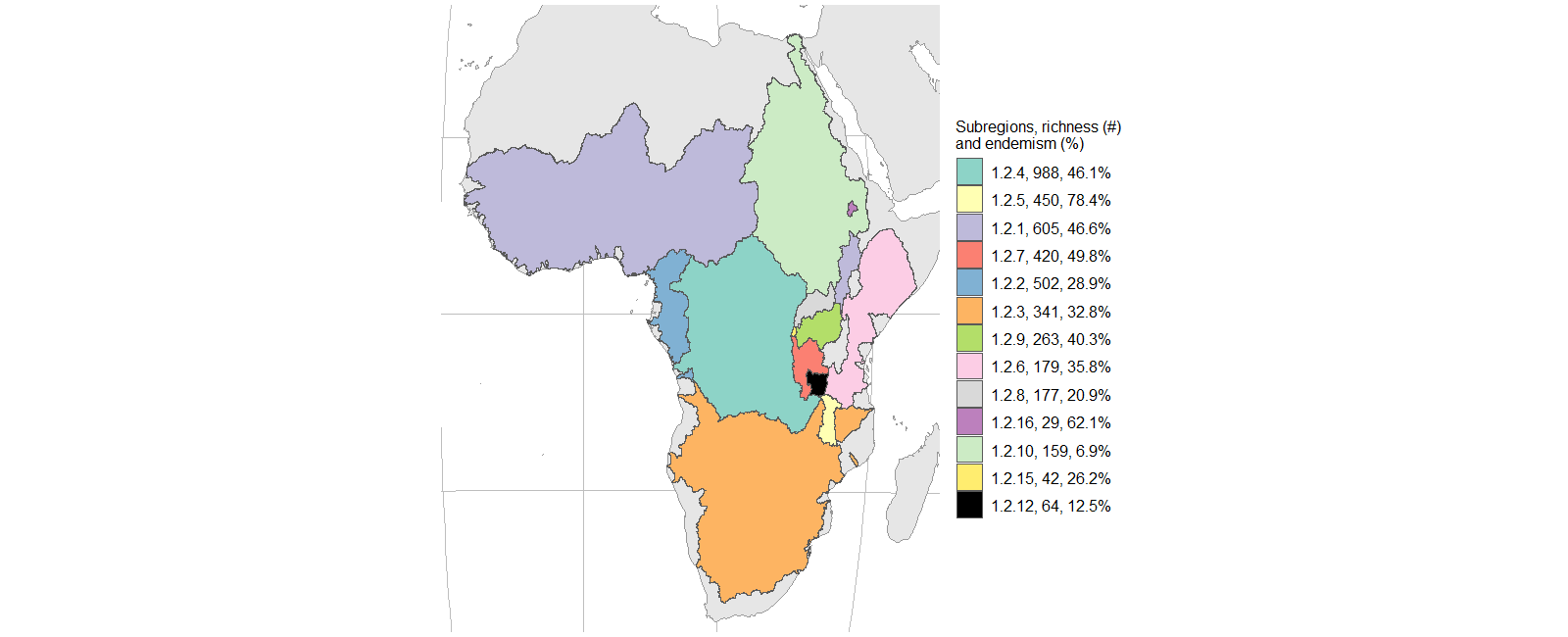

**Figure S7.9.** Subregions of the Afrotropical with species richness and percentage of endemic species. Basins with less than 0.5% of total region richness were removed.

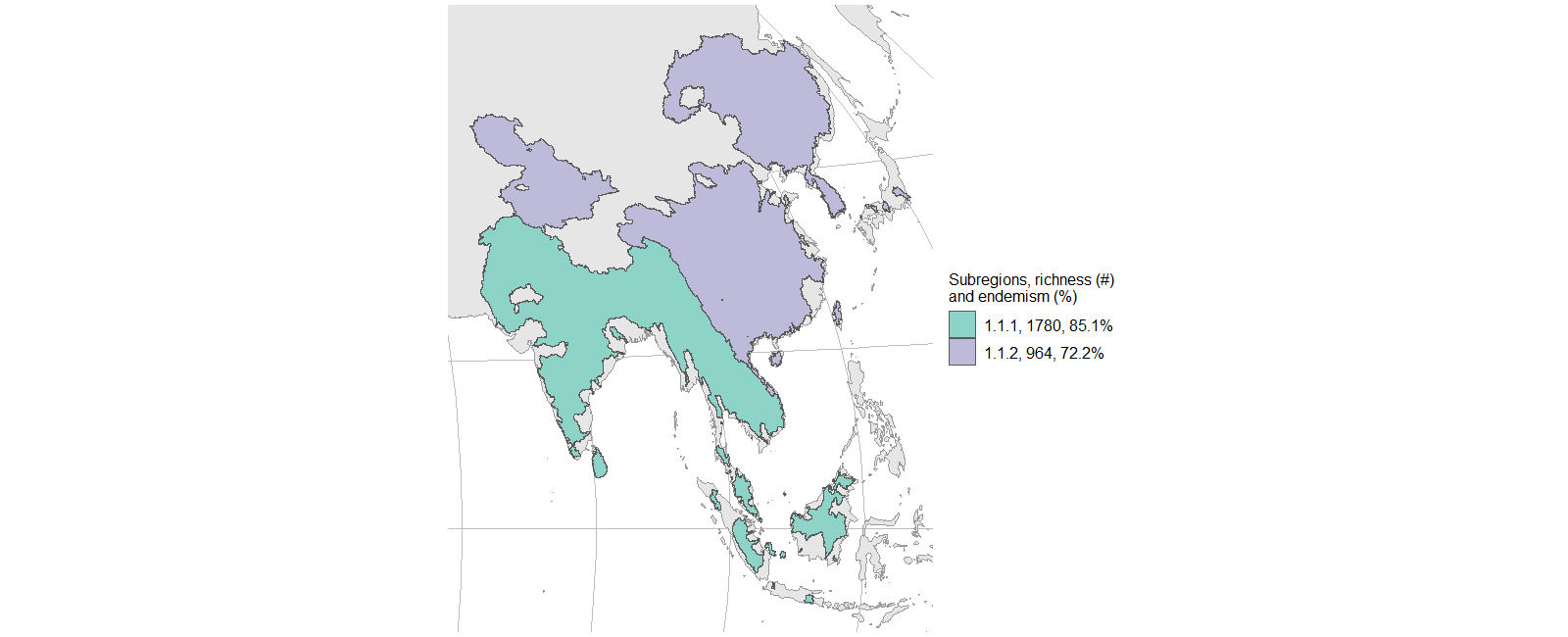

**Figure S7.10.** Subregions of the Sino-Oriental region with species richness and percentage of endemic species. Basins with less than 0.5% of total region richness were removed.

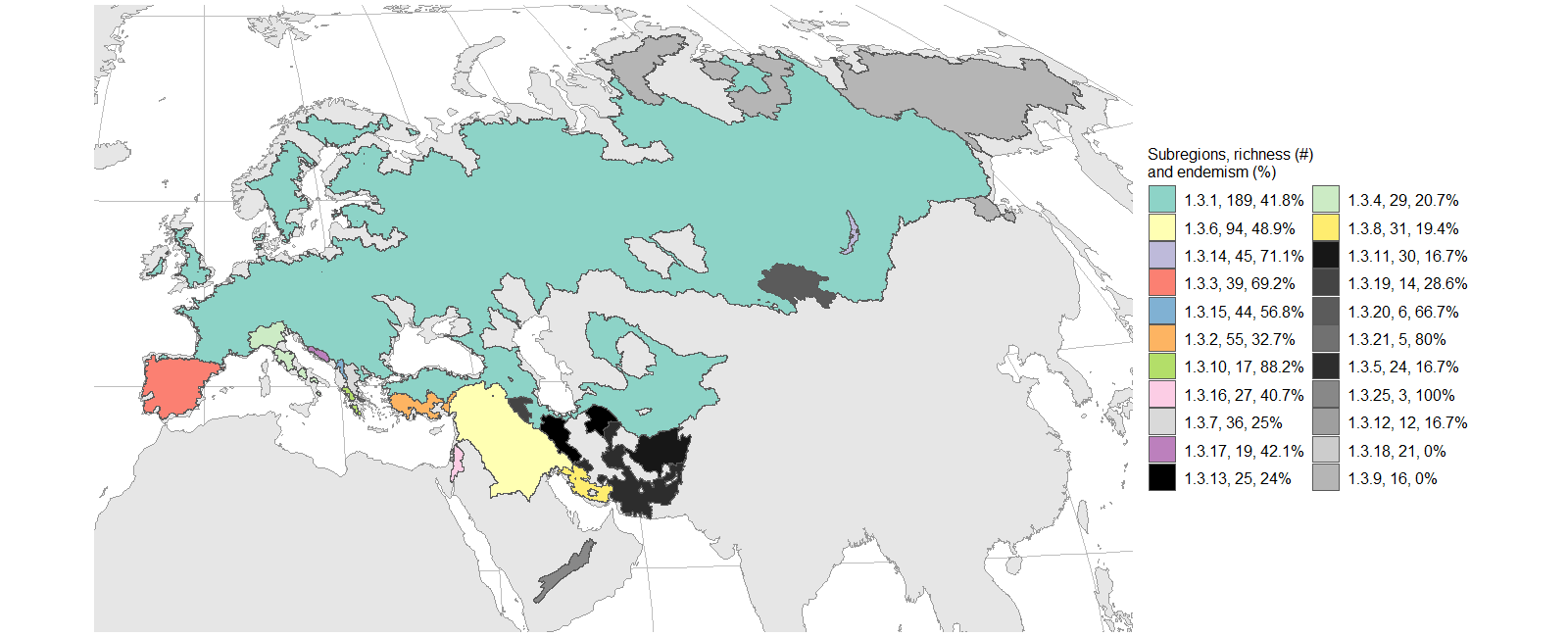

**Figure S7.11.** Subregions of the Palearctic region with species richness and percentage of endemic species. Basins with less than 0.5% of total region richness were removed.

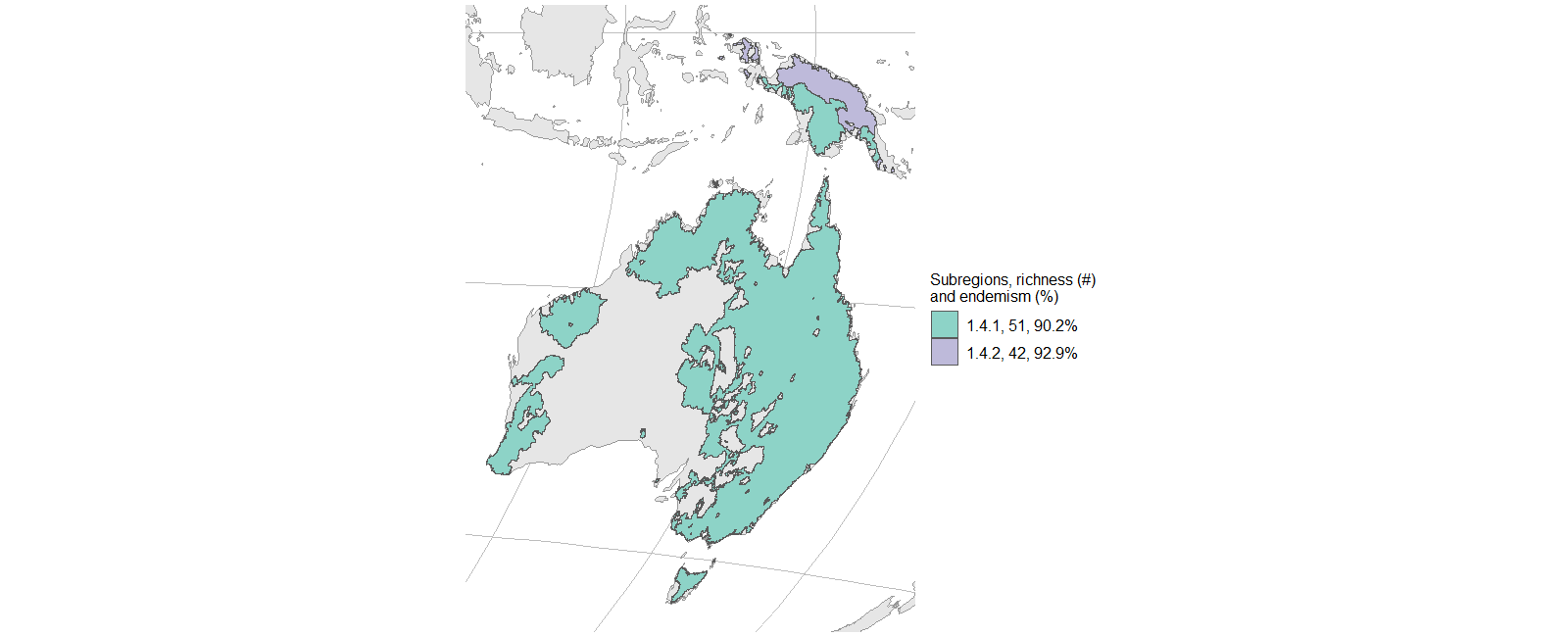

**Figure S7.12.** Subregions of the Oceanian region with species richness and percentage of endemic species. Basins with less than 0.5% of total region richness were removed.

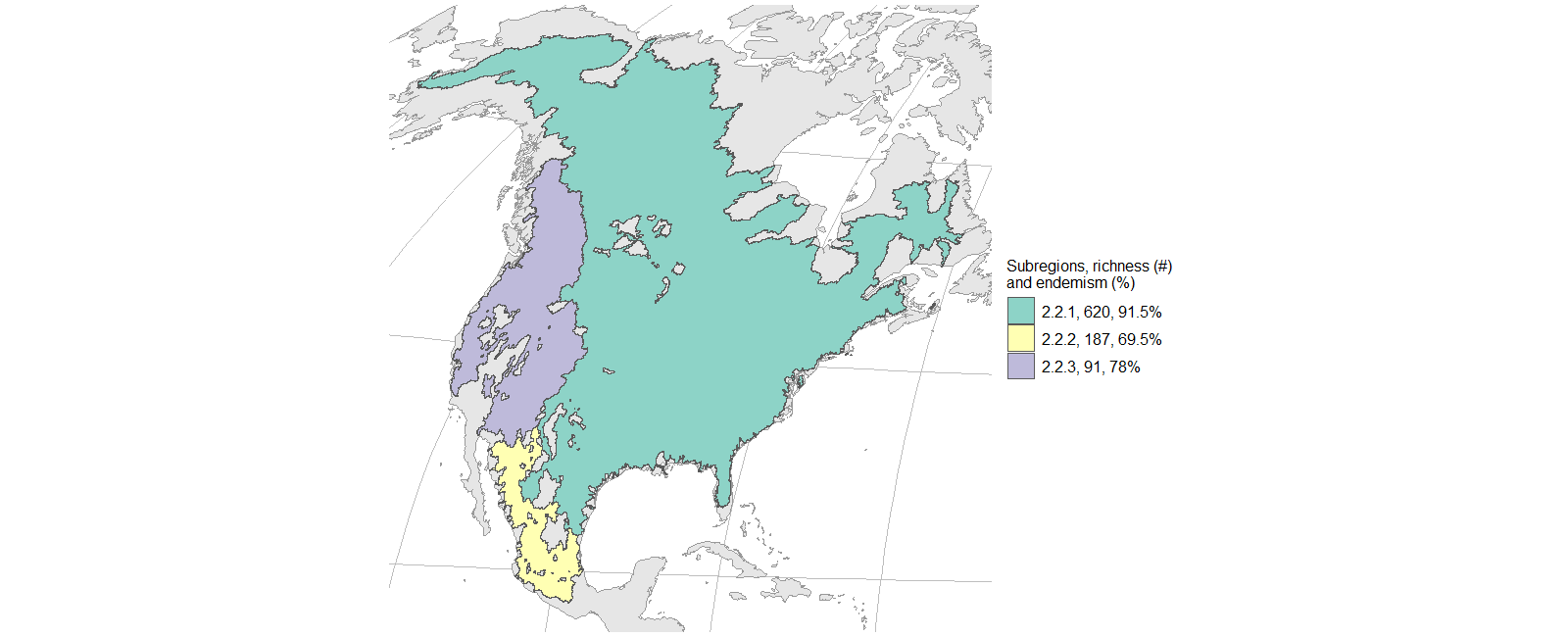

**Figure S7.13.** Subregions of the Nearctic region with species richness and percentage of endemic species. Basins with less than 0.5% of total region richness were removed.

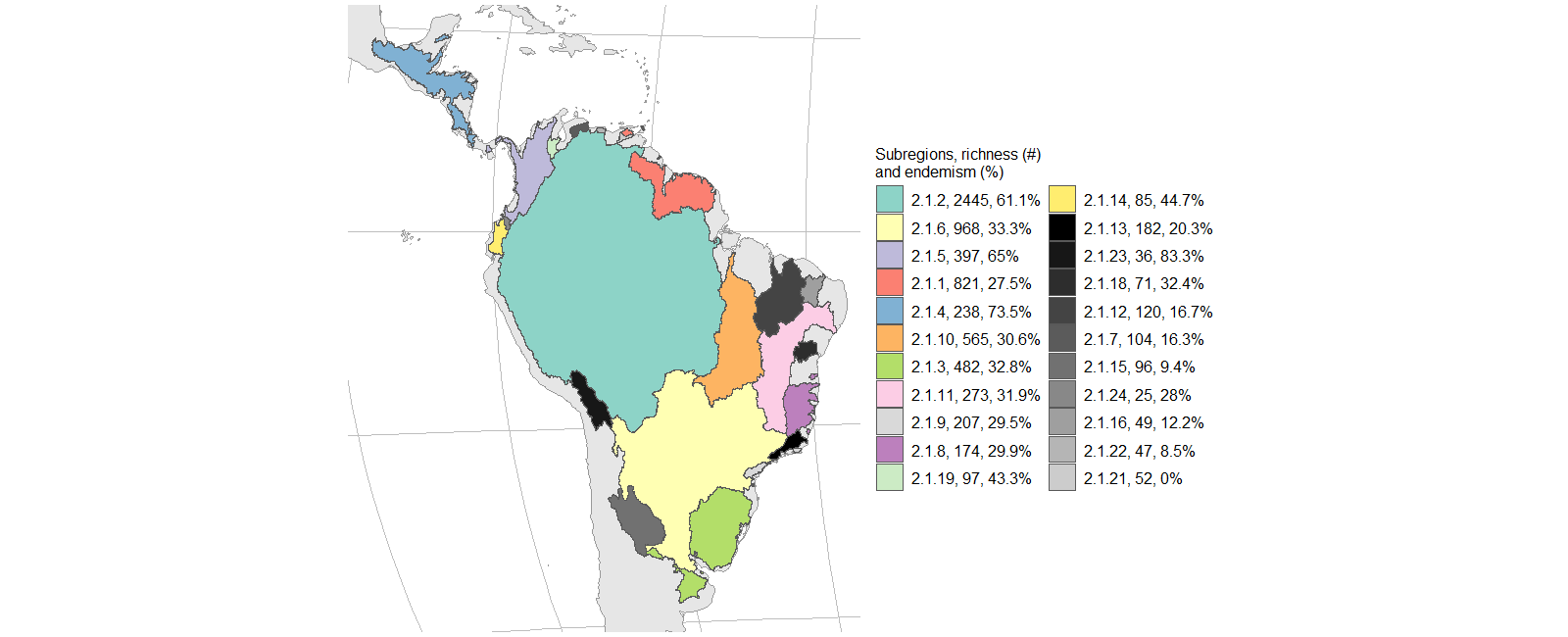

**Figure S7.14.** Subregions of the Neotropical region with species richness and percentage of endemic species. Basins with less than 0.5% of total region richness were removed.

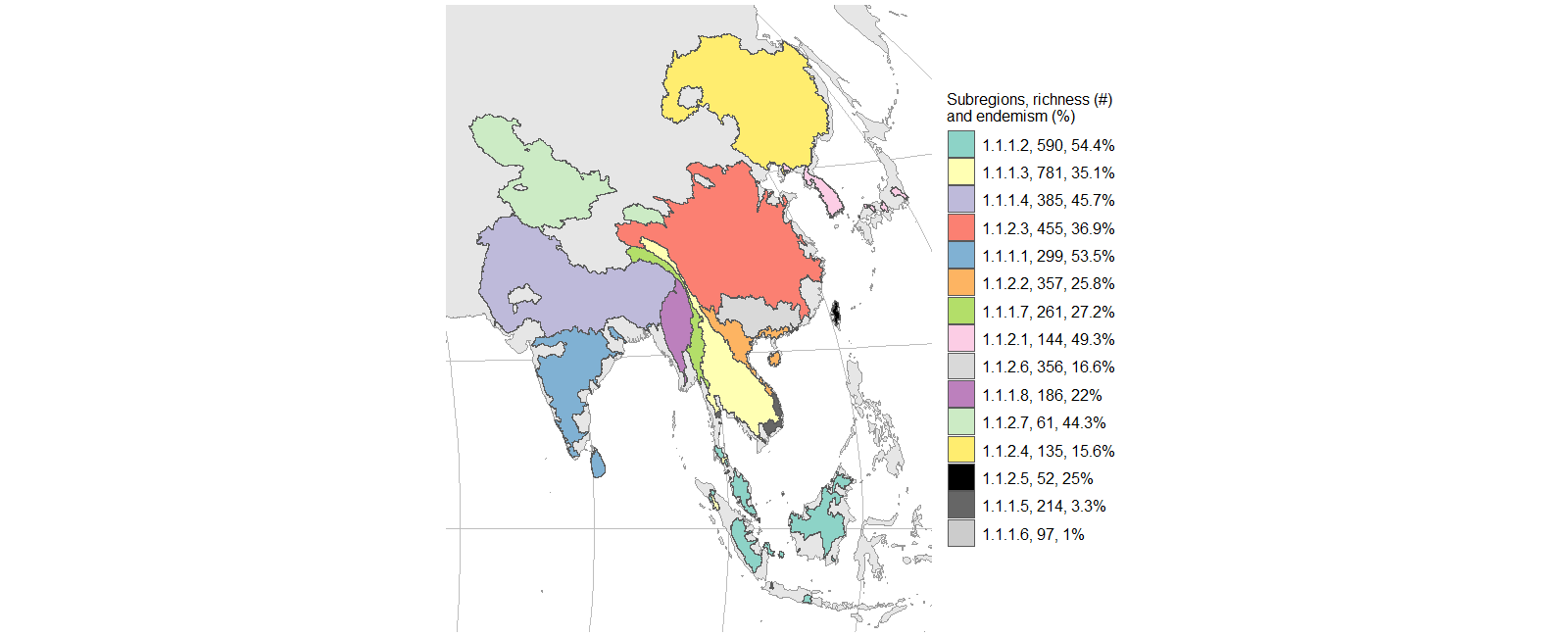

**Figure S7.15.** Level 4 regions of the Sino-Oriental region with species richness and percentage of endemic species. Basins with less than 0.5% of total region richness were removed.
