## Appendix S8 for "Global biogeographical regions of freshwater fish species"

*Preprint manuscript submitted to the Journal of Biogeography*

**Appendix S8. Endemism rates for other vertebrate groups.**

- **Procheş & Ramdhani's World Zoogeographical Regions (2012) of amphibians, mammals and vertebrates**

We worked on the original database of species occurrences in ecoregions as provided by Procheş & Ramdhani. For the results to be comparable with our study, we merged their subregions into major regions according to Figure 2 in Procheş & Ramdhani (2012).

| Table S8.2. Richness and endemism of birds, mammals and herptiles in the major regions of Procheş & Ramdhani (2012). Major regions (i.e., regions similar in size to our regions) were highlighted in bold. | | | | | | | | | | | |
| --- | --- | --- | --- | --- | --- | --- | --- | --- | --- | --- | --- |
|  | Birds | | |  | Mammals | | |  | Herptiles | | |
|  | Number of species | Number of endemic species | Percentage of endemic species |  | Number of species | Number of endemic species | Percentage of endemic species |  | Number of species | Number of endemic species | Percentage of endemic species |
| Regions |  |  |  |  |  |  |  |  |  |  |  |
| **Neotropical** | **3391** | **1875** | **55.3%** |  | **1179** | **543** | **46.1%** |  | **3968** | **3204** | **80.7%** |
| **Ethiopian** | **1955** | **1504** | **76.9%** |  | **1002** | **900** | **89.8%** |  | **1884** | **1781** | **94.5%** |
| **Palearctic** | **1734** | **293** | **16.9%** |  | **1026** | **542** | **52.8%** |  | **1484** | **1093** | **73.7%** |
| **Indo-Malaysian** | **1646** | **445** | **27.0%** |  | **757** | **339** | **44.8%** |  | **1616** | **1228** | **76.0%** |
| Andean | 1252 | 231 | 18.5% |  | 454 | 70 | 15.4% |  | 655 | 362 | 55.3% |
| **Nearctic** | **960** | **109** | **11.4%** |  | **593** | **206** | **34.7%** |  | **851** | **390** | **45.8%** |
| Wallacean | 951 | 451 | 47.4% |  | 340 | 225 | 66.2% |  | 577 | 374 | 64.8% |
| New Guinean | 830 | 484 | 58.3% |  | 268 | 198 | 73.9% |  | 621 | 470 | 75.7% |
| Caribbean | 659 | 166 | 25.2% |  | 94 | 44 | 46.8% |  | 560 | 514 | 91.8% |
| **Australian** | **600** | **306** | **51.0%** |  | **268** | **221** | **82.5%** |  | **989** | **903** | **91.3%** |
| Polynesian | 460 | 250 | 54.3% |  | 35 | 24 | 68.6% |  | 164 | 136 | 82.9% |
| Madagascan | 311 | 189 | 60.8% |  | 148 | 136 | 91.9% |  | 528 | 510 | 96.6% |

- **Holt et al., (2013)’s update** **of Wallace’s zoogeographic regions of the World**

We worked on the list of species names as provided by Holt and colleagues and the 2018-2 version of IUCN distribution data (IUCN 2018) for terrestrial mammals and amphibians (downloaded 15/11/2018), and the birdlife species distribution maps of the world version 7.0 (downloaded 15/11/2018). We checked for synonyms with the 2018-2 version of the IUCN database using the rl_synonyms function of the rredlist R package version 0.5.0. For mammals, we were able to recover 331 synonyms of the 450 species names of Holt et al. (2013) that did not exist in the 2018-2 IUCN datasets. We excluded the remaining unknown 119 species (i.e., 2.5% of Holt et al. species). For amphibians, we were able to recover 651 synonyms of the 689 species names of Holt et al. (2013) that did not exist in the 2018-2 IUCN datasets. We excluded the remaining unknown 38 species (i.e., 0.6% of Holt et al. species). For birds, we were able to recover 954 synonyms of the 1511 species names of Holt et al. (2013) that did not exist in the 2018-2 IUCN datasets. We excluded the remaining unknown 554 species (i.e., 5.5% of Holt et al. species). We included only the native range of species, and, for birds, only the breeding range.

| Table S8.3. Richness and endemism of birds, mammals and herptiles in the major regions of Holt et al. (2013). Major regions (i.e., regions similar in size to our regions) were highlighted in bold. | | | | | | | | | | | |
| --- | --- | --- | --- | --- | --- | --- | --- | --- | --- | --- | --- |
|  | Birds | | |  | Mammals | | |  | Amphibians | | |
|  | Number of species | Number of endemic species | Percentage of endemic species |  | Number of species | Number of endemic species | Percentage of endemic species |  | Number of species | Number of endemic species | Percentage of endemic species |
| Regions |  |  |  |  |  |  |  |  |  |  |  |
| **Neotropical** | **2928** | **1729** | **59.1%** |  | **949** | **622** | **65.5%** |  | **2133** | **1833** | **85.9%** |
| **Oriental** | **1975** | **910** | **46.1%** |  | **1041** | **606** | **58.2%** |  | **941** | **761** | **80.9%** |
| Panamanian | 1756 | 323 | 18.4% |  | 566 | 142 | 25.1% |  | 1051 | 695 | 66.1% |
| **Afrotropical** | **1732** | **1453** | **83.9%** |  | **983** | **849** | **86.4%** |  | **720** | **707** | **98.2%** |
| **Oceanina** | **1129** | **733** | **64.9%** |  | **386** | **252** | **65.3%** |  | **392** | **363** | **92.6%** |
| Sino-Japanese | 1102 | 20 | 1.8% |  | 511 | 49 | 9.6% |  | 343 | 155 | 45.2% |
| **Palearctic** | **1060** | **125** | **11.8%** |  | **619** | **190** | **30.7%** |  | **154** | **101** | **65.6%** |
| **Nearctic** | **998** | **282** | **28.3%** |  | **549** | **316** | **57.6%** |  | **431** | **363** | **84.2%** |
| Saharo-Arabian | 834 | 21 | 2.5% |  | 394 | 41 | 10.4% |  | 66 | 15 | 22.7% |
| **Australian** | **649** | **346** | **53.3%** |  | **261** | **179** | **68.6%** |  | **224** | **198** | **88.4%** |
| Madagascan | 220 | 150 | 68.2% |  | 132 | 121 | 91.7% |  | 234 | 233 | 99.6% |

BirdLife International and Handbook of the Birds of the World (2017) Bird species distribution maps of the world. Version 2017.2. Available at <http://datazone.birdlife.org/species/requestdis>. Downloaded on 15 November 2018.

Holt, B. G., Lessard, J.-P., Borregaard, M. K., Fritz, S. A., Araújo, M. B., Dimitrov, D., … Rahbek, C. (2013). An update of Wallace’s zoogeographic regions of the world. *Science*, *339*(6115), 74–78. http://doi.org/10.1126/science.1228282

IUCN 2018. *The IUCN Red List of Threatened Species. Version 2018-2* <[http://www.iucnredlist.org](http://www.iucnredlist.org/)>. Downloaded on 15 November 2018.

Procheş, Ş., & Ramdhani, S. (2012). The World’s Zoogeographical Regions Confirmed by Cross-Taxon Analyses. *BioScience*, *62*(3), 260–270. http://doi.org/10.1525/bio.2012.62.3.7
